# Confidence-aware learning for transcriptome-based prediction of OXPHOS genes in *Caenorhabditis elegans* under incomplete functional annotation

**DOI:** 10.64898/2026.05.29.728242

**Authors:** Sofía Zeballos-Gorón, Gustavo Salinas, Flavio Pazos Obregón

## Abstract

Assigning biological functions to genes remains a major challenge in genomics because reliable functional annotations are often scarce and unevenly distributed. Although transcriptomic datasets capture rich information about gene activity and coordination, exploiting these data for gene function prediction is difficult when only a small number of genes have high-confidence functional assignments and the remaining labels are uncertain.

Here, we developed a confidence-aware learning framework that integrates complementary transcriptomic landscapes while explicitly incorporating uncertainty in functional annotations. The framework combines supervised learning from time-resolved bulk RNA-seq data with co-expression analysis of embryonic and adult single-cell transcriptomes. Supervised learning uses a two-round training scheme in which genes supported by limited functional evidence are incorporated only after an initial model has been established using high-confidence annotations. To reduce module-specific biases, we implemented an informed bagging strategy in which genes from individual functional modules are systematically withheld during training and predictions are integrated by consensus across models.

We applied this framework to identify genes involved in oxidative phosphorylation (OXPHOS) in *Caenorhabditis elegans*. Integrating supervised and co-expression evidence prioritized a small set of high-confidence candidate genes with strong predictive performance on an independent test set. Experimental validation showed that disruption of the top-ranked candidate, *ril-1*, produces phenotypes consistent with impaired OXPHOS function.

Our results demonstrate that transcriptomic landscapes can be systematically exploited for gene function prediction under incomplete functional annotation, providing a framework that may be transferable to other biological processes and organisms where reliable annotations remain limited.

## Introduction

Understanding gene function remains one of the central challenges in biology. Despite decades of experimental and computational efforts, a substantial fraction of genes across sequenced genomes still lack reliable functional annotation, limiting our ability to fully understand cellular processes and biological systems (1,2). Traditional approaches based on sequence homology have been instrumental for transferring functional information between organisms, but they face intrinsic limitations: many genes lack close homologs, whereas paralogous genes may retain high sequence similarity despite functional divergence (3,4). More recently, structure-based annotation has expanded the reach of homology-based prediction (5,6), yet structural similarity alone does not necessarily imply functional equivalence, and proteins with distinct folds may converge on similar biological roles. Consequently, complementary strategies capable of exploiting orthogonal sources of biological information are increasingly needed.

Large-scale transcriptomic datasets constitute one such source of information. Genes participating in the same biological process often exhibit coordinated expression across developmental stages, tissues, or physiological conditions, providing functional signals that are largely independent of sequence conservation (7,8). Recent advances in machine learning have enabled increasingly accurate exploitation of transcriptomic data for gene function prediction by integrating heterogeneous biological information in a systematic and scalable manner (9,10). However, a fundamental challenge remains largely overlooked. Supervised learning relies on annotated genes as training labels, yet biological annotations differ markedly in both completeness and confidence. In practice, only a relatively small subset of genes can be considered high-confidence positives, whereas many others are supported by limited or indirect evidence. Consequently, gene function prediction is constrained not only by the availability of transcriptomic data but also by the scarcity and uneven confidence of functional annotations. From a machine learning perspective, this problem shares characteristics with weakly supervised and positive-unlabeled learning, where only a subset of positive instances can be regarded as reliable training examples while the remaining data carry uncertain or incomplete labels (11–13). Although these learning paradigms have been extensively investigated in computer science, they have been only sparsely explored in transcriptome-based gene function prediction, where annotation confidence itself may provide valuable information for model construction.

Oxidative phosphorylation (OXPHOS) provides an excellent system for addressing this challenge. OXPHOS is the central bioenergetic pathway in mitochondria that couples electron transfer through the electron transport chain (ETC) to ATP synthesis. The ETC comprises four multi-subunit protein complexes: I (NADH dehydrogenase), II (succinate dehydrogenase), III (cytochrome c reductase), and IV (cytochrome c oxidase), embedded in the inner mitochondrial membrane, whichtransfer electrons from the reduced substrates NADH and succinate to molecular oxygen. The energy released during electron flow is harnessed to pump protons from the mitochondrial matrix into the intermembrane space, generating an electrochemical proton gradient, which drives ATP synthesis through complex V (ATP synthase), thus completing the process of OXPHOS (14–16).

The OXPHOS complexes are composed of both core and accessory proteins. Core subunits, encoded by the mitochondrial and nuclear genomes, are essential for the primary catalytic function of the complexes. The accessory subunits, though non-catalytic, stabilize complexes and modulate their activity. These subunits are often less conserved evolutionarily, and their number can vary significantly among species, potentially underlying lineage-specific adaptations. Depending on the species, the OXPHOS complexes comprise 80-100 protein subunits. In addition, several assembly factors are required for the proper folding, maturation, cofactor incorporation, and stepwise assembly of the complexes from their individual subunits, although these factors are not part of the final functional complexes. While the core subunits of OXPHOS are well characterized in mammals, some regulatory and accessory subunits, as well as assembly factors remain unidentified, particularly in non-mammalian lineages. The dynamic formation of supercomplexes (respirasomes) and tissue-specific regulation further contribute to the complexity of this system (17–19).

Identifying the complete set of OXPHOS-associated genes is important not only for understanding mitochondrial biology but also because defects in OXPHOS are among the most common causes of mitochondrial diseases in humans, a heterogeneous group of metabolic disorders that primarily affect tissues with high energy demand such as the brain, muscle, heart, and liver. Pathogenic mutations in genes encoding complex subunits or assembly factors lead to impaired electron transfer, reduced ATP production, and excessive generation of reactive oxygen species. Clinically, OXPHOS deficiencies, most often lethal, manifest in a wide range of syndromes, including Leigh syndrome, Leber hereditary optic neuropathy, mitochondrial encephalomyopathy, myopathy, cardiomyopathy, and neurodegeneration (20,21).

*Caenorhabditis elegans* represents an ideal model organism in which to address this problem. Approximately 60–80% of human disease genes have recognizable orthologs in *C. elegans*, and many essential cellular pathways, including mitochondrial metabolism, are evolutionarily conserved (22,23). Extensive and high-quality transcriptome resources are available for *C. elegans*: bulk and single-cell RNA-seq datasets allow multi-resolution exploration of gene expression across tissues, developmental stages, and environmental conditions (24,25).

The conservation of OXPHOS machinery makes *C. elegans* an excellent in vivo system for modeling human mitochondrial diseases and to dissect how OXPHOS influences development, stress responses, and aging. Partial defects in OXPHOS genes in *C. elegans* produce a variety of well-characterized phenotypes, including slowed development, reduced fertility, altered locomotion, and changes in stress resistance and lifespan. Widely studied mutants such as *isp-1* (complex III), *mev-1* (complex II), *nuo-6* and *gas-1* (complex I) display impaired respiration and mimic aspects of human mitochondrial dysfunction (26–28). Remarkably, mild reductions in respiratory activity can trigger adaptive stress responses, including activation of the mitochondrial unfolded protein response (UPRmt) and ROS-dependent signaling pathways that extend lifespan, illustrating a conserved mitohormetic mechanism (29,30).

In addition to the relevance of *C. elegans* to understand and model human diseases, *C. elegans* is a free-living nematode widely used as a genetic and biochemical model for parasitic nematodes (31). Parasitic worms infect humans, livestock, companion animals, and crops, representing a major burden for health and agriculture worldwide. Notably, both *C. elegans* and parasitic nematodes possess an alternative electron transport chain (aETC) that operates under hypoxic conditions and relies on rhodoquinone-dependent electron transport and fumarate reduction (32–34). In addition, several parasitic helminths exhibit lineage-specific duplications of ETC-related genes that may contribute to metabolic adaptation to fluctuating oxygen availability (35,36). Importantly, mitochondrial respiratory complexes, particularly complex II and rhodoquinone-dependent metabolism, have emerged as promising pharmacological targets against parasitic nematodes (37).

Here, we addressed gene function prediction under incomplete functional annotation by developing a confidence-aware learning strategy that integrates complementary transcriptomic landscapes while explicitly accounting for different levels of confidence in existing biological knowledge. Our framework combines supervised learning on time-resolved bulk RNA-seq data with functional inference from embryonic and adult single-cell co-expression networks. To address incomplete functional annotations, supervised learning incorporated a two-round learning strategy in which genes supported by limited functional evidence were introduced only after an initial model had been established using high-confidence annotations. To reduce module-specific biases within the positive training set, we further implemented a novel informed bagging procedure in which genes belonging to individual functional modules were iteratively withheld during training, and consensus predictions were obtained across the resulting models. We applied this framework to identify previously unrecognized genes involved in oxidative phosphorylation in *C. elegans* and experimentally validated one of the highest-ranked candidates. More broadly, this work explores how transcriptomic landscapes and confidence-aware learning can be combined to improve gene function prediction in biological systems where reliable functional annotations remain limited.

## 2 Results

The overall confidence-aware learning framework developed in this study is summarized in Figure 1.

**Figure 1.**
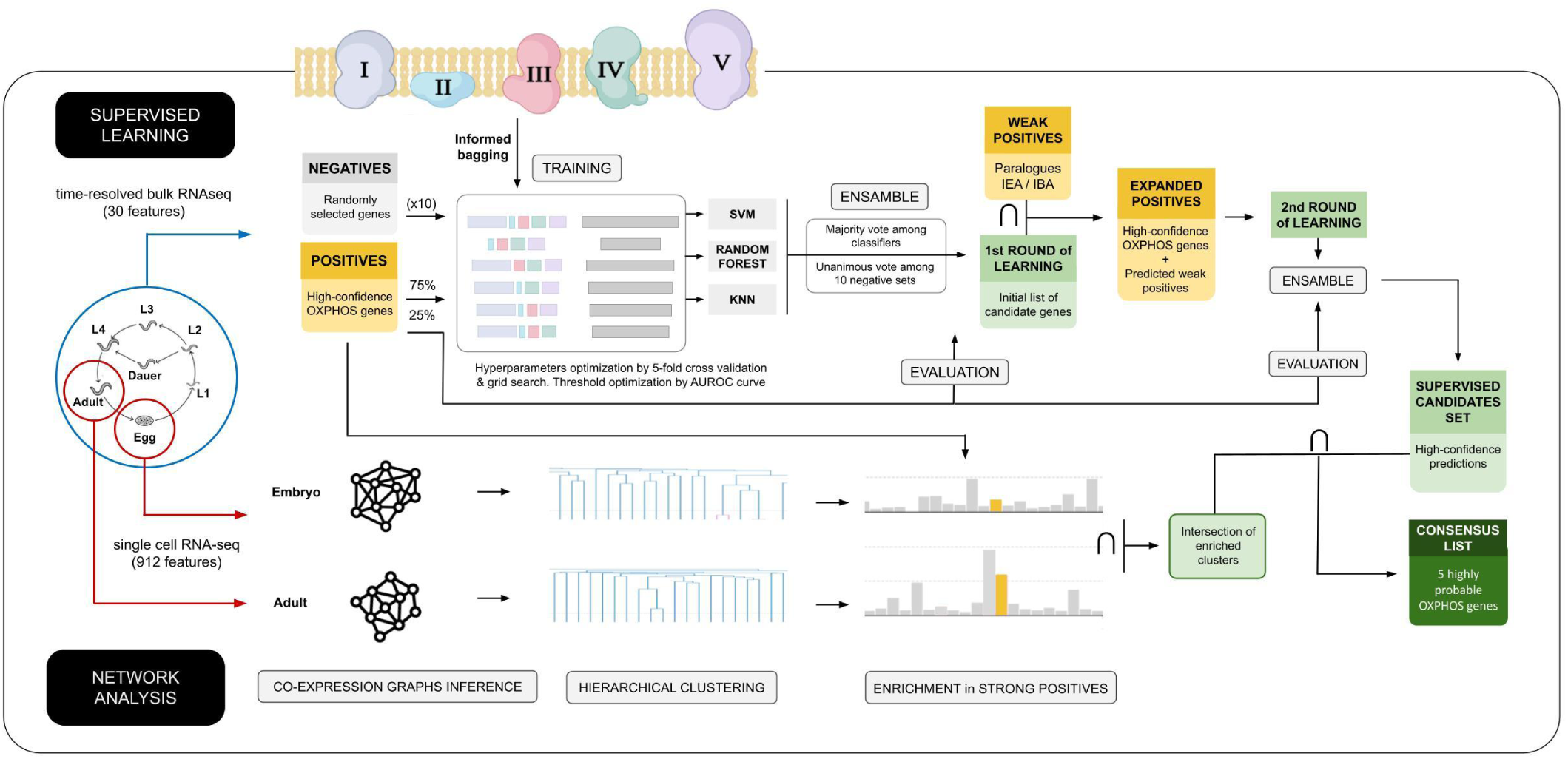
Confidence-aware learning framework for transcriptome-based OXPHOS gene prediction in C. elegans. The upper panel shows the supervised component of the framework. Time-resolved bulk RNA-seq data spanning embryonic, larval, dauer, and adult stages were combined with a curated set of high-confidence OXPHOS genes and ten independently generated negative sets to train support vector machine (SVM), random forest (RF), and k-nearest neighbors (KNN) classifiers. To reduce module-specific biases within the positive training set, an informed bagging strategy was implemented in which five positive training configurations were generated by systematically excluding one OXPHOS complex at a time, together with an additional configuration containing genes from all complexes. Combined with the ten negative sets, this strategy produced 180 binary classifiers. Predictions were integrated through a two-level ensemble voting procedure to generate high-confidence candidates from the first learning round. Low-confidence positives recovered with high confidence were then incorporated into an expanded positive training set for a second learning round, yielding the final supervised candidate set.

The lower panel shows the co-expression component of the framework. Independent embryonic and adult single-cell RNA-seq datasets were used to construct gene co-expression networks, followed by hierarchical clustering and identification of OXPHOS-enriched modules. Genes belonging to enriched modules in both developmental contexts were retained as unsupervised candidates. Finally, the supervised candidate set was intersected with the OXPHOS-enriched modules from both embryonic and adult networks to generate the final consensus list of high-confidence OXPHOS candidate genes.

### 2.1 Supervised approach

#### High-confidence OXPHOS genes

We assembled a high-confidence list of genes for which there is evidence of their participation in oxidative phosphorylation in *C. elegans (see Methods)*. This integrative curation resulted in a list of 64 genes used as positive examples for training and evaluation (Table S1).

#### Low-confidence positives, including paralogs

Besides the high-confidence positives, we assembled a second list of genes supported only by indirect evidence (GO IEA/IBA annotations or unresolved paralogs). Rather than using these genes during initial training, we treated them as low-confidence positives and used them to evaluate whether the models could recover biologically plausible candidates. Table S2 shows the complete list of low confidence positives.

#### First round of learning

We first evaluated whether the confidence-aware learning framework could accurately recognize canonical OXPHOS genes using only the high-confidence set for training. To this end, 25% of the high-confidence positives were withheld as an independent evaluation set, while the remaining genes were used to train ensembles of Random Forest, Support Vector Machine, and k-Nearest Neighbor classifiers following the informed bagging strategy (see Methods).

All models showed high performance metrics over the independent evaluation set despite being trained on multiple alternative negative sets (Table 1). Mean F1-scores reached 0.94 across classifiers trained on the Boeck *et al.* transcriptomic compendium, demonstrating that the framework robustly recovered held-out OXPHOS genes. These results indicate that the informed bagging strategy yields stable models even in the absence of a reliable negative training set. External generalization to independent bulk transcriptomic datasets was not evaluated.

**Table 1-.**
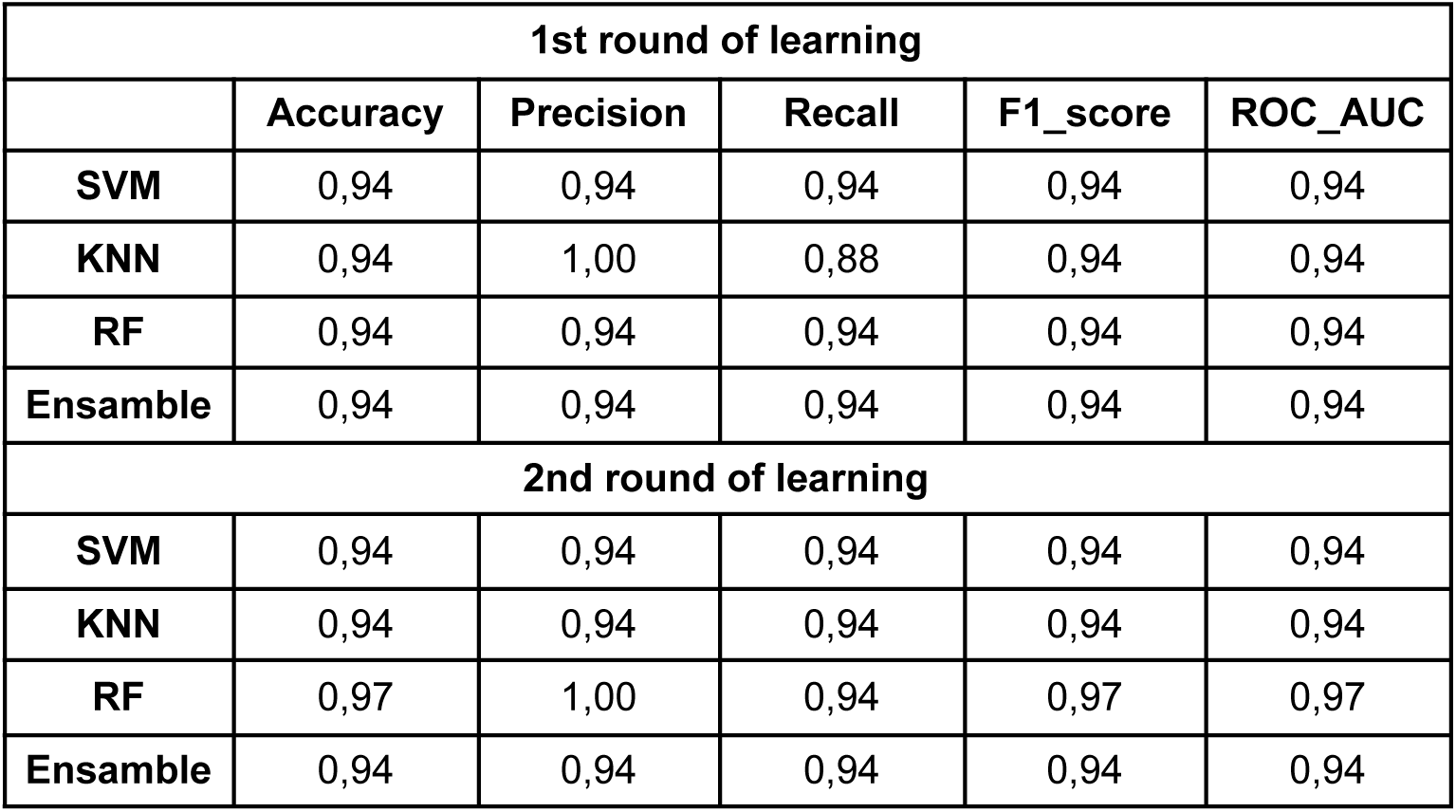
Performance of classifiers.

| 1st round of learning |  |  |  |  |  |
| --- | --- | --- | --- | --- | --- |
|  | Accuracy | Precision | Recall | F1_score | ROC_AUC |
| **SVM** | 0,94 | 0,94 | 0,94 | 0,94 | 0,94 |
| **KNN** | 0,94 | 1,00 | 0,88 | 0,94 | 0,94 |
| **RF** | 0,94 | 0,94 | 0,94 | 0,94 | 0,94 |
| **Ensamble** | 0,94 | 0,94 | 0,94 | 0,94 | 0,94 |
| 2nd round of learning |  |  |  |  |  |
| **SVM** | 0,94 | 0,94 | 0,94 | 0,94 | 0,94 |
| **KNN** | 0,94 | 0,94 | 0,94 | 0,94 | 0,94 |
| **RF** | 0,97 | 1,00 | 0,94 | 0,97 | 0,97 |
| **Ensamble** | 0,94 | 0,94 | 0,94 | 0,94 | 0,94 |

Once trained, each model was used to classify the remaining *C. elegans* genes based on their expression patterns. The number of genes predicted as positive varied substantially across models, and after applying the ensemble rules detailed in Methods we obtained a list of genes classified as OXPHOS genes by our classifiers after the first round of learning.

#### Recovery of low-confidence positives and paralog discrimination

To evaluate whether the framework generalized beyond the curated training set, we examined the classification of genes that had been intentionally excluded from training because of low-confidence functional evidence. We considered as recovered only genes predicted with a mean probability greater than 0.9 across the ensemble of first-round classifiers.

Among the low-confidence genes supported only by indirect evidence (GO IEA/IBA annotations), 6 of 8 genes were recovered with high confidence, including candidates associated with complexes I, III, and IV (Table S2). Thus, genes supported only by indirect GO annotations or limited prior evidence were frequently classified as OXPHOS components despite not contributing to model training.

An interesting pattern emerged for duplicated and triplicated OXPHOS paralogs. Rather than predicting all members of a paralogous family, the models generally assigned high confidence to a single representative copy. High-confidence predictions were obtained for *ndab-2, ucr-1, cox-6c, cyc-2.1, asb-2, asg-2,* and *hpo-18*, whereas several other paralog groups remained below the probability threshold (Table S2). Notably, some excluded paralogs received probabilities close to the cutoff (for example, *nduv-3, ndab-1, F45H10.2, R07E4.3,* and *asg-1*), whereas others were assigned very low probabilities (less than 0.2), suggesting that the framework distinguished between genes with transcriptional profiles compatible with canonical OXPHOS function and genes unlikely to belong to the core OXPHOS program. Overall, 7 of the 25 duplicated or triplicated paralog genes were recovered with high confidence, consistent with a preferential selection of a single representative copy within most paralog groups.

#### Assembly factors provide a biological specificity control

To further assess the biological specificity of the learned transcriptional signature, we evaluated genes annotated as respiratory complex assembly factors. None of the 34 assembly factor genes (Table S3) was classified as positive by the first-round ensemble. Because these proteins participate in transient assembly processes rather than as components of the mature respiratory complexes, their systematic exclusion provides an independent biological validation that the framework preferentially captures the transcriptional program of core and accessory OXPHOS components.

#### Second round of confidence-aware learning

The low-confidence positives recovered with high confidence in the first learning round were subsequently used to expand the positive training set, implementing the confidence-aware learning strategy illustrated in Figure 1. Incorporation of the recovered low-confidence positives increased the number of genes used for training from 48 to 61 genes. We then repeated the complete training and ensembling procedure and evaluated the resulting models using the same test set as in the first learning round. The corresponding performance metrics are reported in Table 1. We subsequently intersected the predictions obtained in the first and second learning rounds. This conservative strategy yielded 640 reproducibly predicted genes, of which 155 showed a mean prediction probability greater than 0.9. These genes were retained as the high-confidence supervised candidate set used in all subsequent analyses (Table S3).

#### Phenotype enrichment analysis supports the biological relevance of the supervised candidate set

Many of the 155 genes in the supervised candidate set lacked informative Gene Ontology or KEGG annotations, limiting the usefulness of conventional functional enrichment analyses. Rather than treating this as a limitation of the predictions, we took advantage of the extensive phenotype annotations generated by large-scale RNAi studies in *C. elegans*, which provide broad genome-wide gene–phenotype associations through the WormBase phenotype ontology (38).

To assess whether the supervised candidates were associated with phenotypes characteristic of OXPHOS dysfunction, we first identified the phenotype terms most strongly enriched among the high-confidence OXPHOS genes used in the second learning round. We then tested whether these same phenotype categories were enriched in the supervised candidate set. The resulting enrichment profile (Figure 2) revealed strong overrepresentation of phenotypes associated with developmental progression and organismal fitness, including embryonic lethality, larval arrest, slow growth, defective early embryonic development, and extended life span.

**Figure 2:**
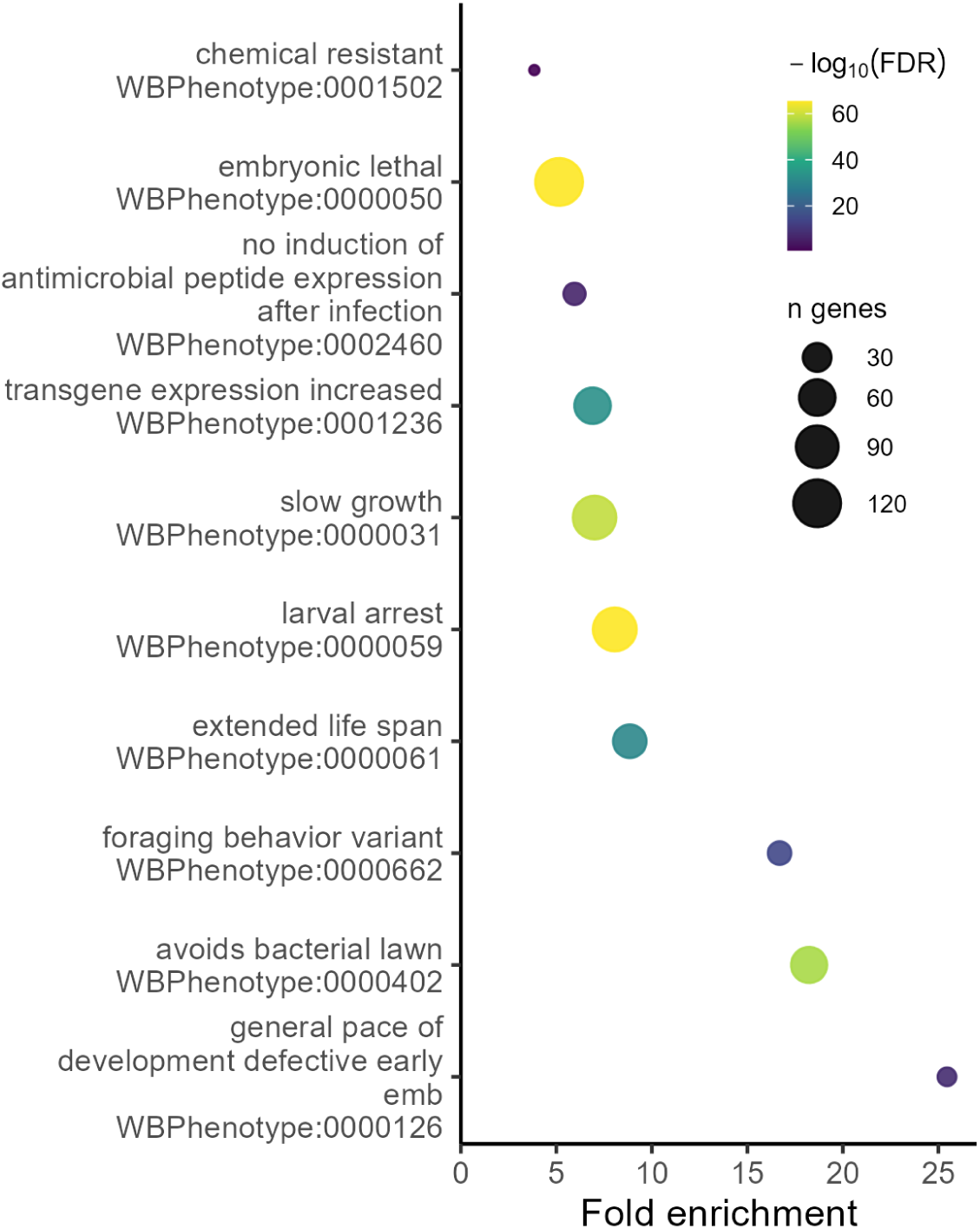
Phenotype enrichment analysis of the supervised candidate set. Phenotype enrichment was evaluated using WormBase RNAi-associated phenotype annotations. The analysis focused on the ten phenotype categories most significantly enriched among the high-confidence OXPHOS genes used in the second learning round and tested whether these same categories were enriched in the supervised candidate set. Dot size indicates the number of genes contributing to each phenotype category, and dot color represents enrichment significance (−log10 FDR), with darker colors indicating stronger enrichment. The enriched phenotype profile is consistent with developmental and physiological defects typically associated with impaired mitochondrial respiration

These phenotypes are consistent with the central role of oxidative phosphorylation in energy production during development, growth, and longevity. The enrichment of lifespan-related phenotypes is particularly noteworthy given the well-established relationship between respiratory chain activity, oxidative phosphorylation, and lifespan regulation in *C. elegans*. Overall, this analysis provides independent biological support for the functional relevance of the supervised candidate set and shows that, despite the limited GO and KEGG annotation of many predicted genes, WormBase phenotype data recover coherent functional signals characteristic of OXPHOS-associated genes and their physiological consequences.

### 2.2 Co-expression network analysis

#### Construction of co-expression networks and mapping of the high-confidence OXPHOS genes and the supervised candidate set

To obtain an independent transcriptome-based view of OXPHOS-associated genes, we constructed co-expression networks from embryonic and adult *C. elegans* single-cell RNA-seq datasets generated by Packer *et al.* and Ghaddar *et al.*, respectively. Hierarchical clustering of each network revealed a single module significantly enriched for the high-confidence OXPHOS genes used in the second learning round (Figure 3, Table S6), indicating that OXPHOS components form a coherent transcriptional program in both developmental contexts.

**Figure 3:**
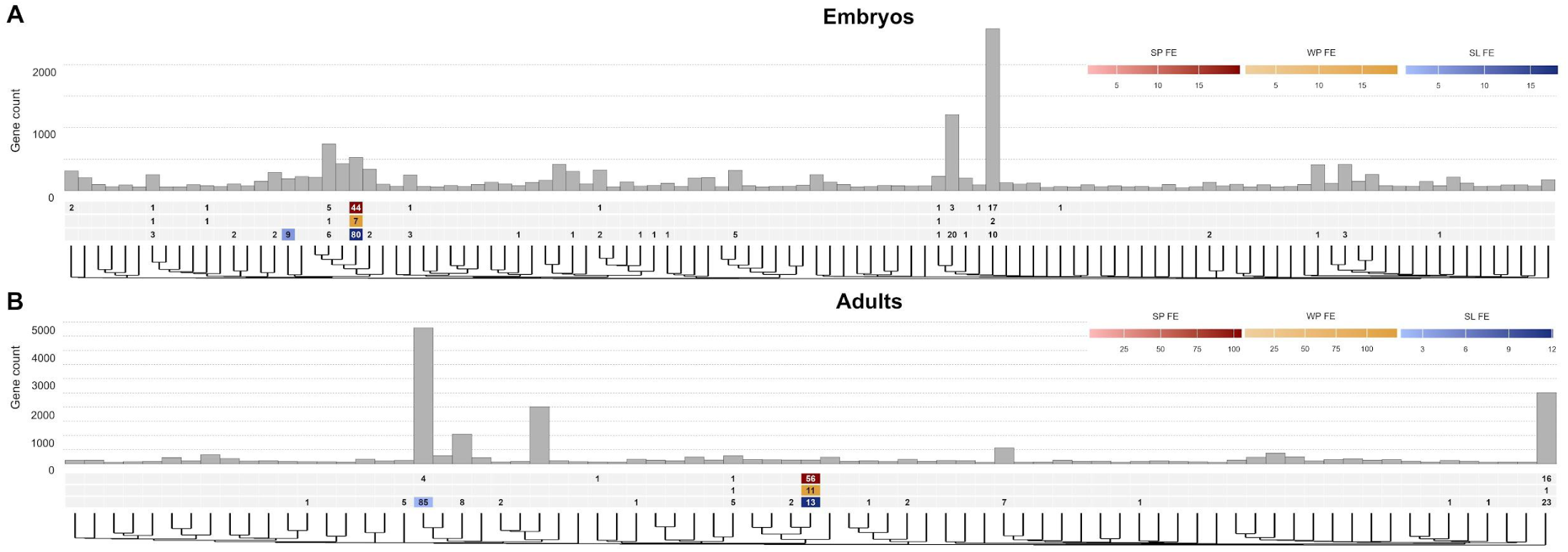
Co-expression network analysis identifies OXPHOS-enriched modules in embryonic and adult single-cell transcriptomes. (A) Hierarchical clustering of the embryonic co-expression network. The dendrogram shows the modules identified from the embryonic single-cell RNA-seq dataset, and bar height indicates the number of genes assigned to each module. The heatmaps below summarize, for each module, the number of high-confidence OXPHOS genes used in the second learning round (red), low-confidence positives recovered after the first round of supervised learning (yellow), and genes belonging to the supervised candidate set (blue). Color intensity represents fold enrichment relative to the genome-wide background, with darker colors indicating stronger enrichment. (B) Hierarchical clustering of the adult co-expression network. Dendrograms, module sizes, heatmaps, and fold-enrichment color scales are presented as described for panel A.

The enrichment signal was substantially stronger in the adult co-expression network than in the embryonic network, with fold enrichments of 105 and 20, respectively (Fisher’s exact test; see Methods). A similar pattern was observed when considering only the low-confidence positives recovered during the first round of supervised learning, which showed fold enrichments of 124 in the adult network and 18 in the embryonic network. Together, these results indicate that adult cell-type expression profiles better resolve the coordinated transcriptional structure associated with OXPHOS-related genes than embryonic profiles under our network construction strategy.

### 2.3 Consensus candidates supported by supervised learning and co-expression analysis

Although the adult co-expression network showed a stronger enrichment signal than the embryonic network, we adopted a conservative integration strategy to prioritize only the most robust OXPHOS candidates. Specifically, we intersected three independent sources of evidence: the supervised candidate set, the OXPHOS-enriched module derived from the adult single-cell transcriptomic dataset, and the OXPHOS-enriched module derived from the embryonic single-cell transcriptomic dataset. This consensus approach retained only genes supported by the confidence-aware supervised framework and independently recovered in OXPHOS-associated co-expression modules across two independent transcriptomic datasets representing distinct developmental contexts.

The intersection yielded a final set of five genes, hereafter referred to as the consensus list: *mai-2, ant-1.1, har-1, ril-1,* and *gmpr-1* (Table 2). These genes represent the highest-confidence candidates identified by our integrative framework and include genes with established mitochondrial functions as well as poorly characterized candidates lacking direct functional evidence. The biological significance of these candidates is examined in the Discussion section, whereas the following section provides experimental evidence supporting the involvement of *ril-1* in OXPHOS.

**Table 2-.** Consensus candidates supported by supervised learning and co-expression analysis.

| Gene | Public Name | Sequence name | Human orthologs | Gene Ontology Association | Automated Description |
| --- | --- | --- | --- | --- | --- |
| WBGene00006439 | *ant-1.1* | T27E9.1 | SLC25A4, SLC25A5, SLC25A6 | "antiporter activity", "ATP:ADP antiporter activity", "determination of adult lifespan", "membrane", "mitochondrial ADP transmembrane transport", "mitochondrial ATP transmembrane transport", "mitochondrial inner membrane", "mitochondrion organization", "mitochondrion", "negative regulation of mitochondrial outer membrane permeabilization involved in apoptotic signaling pathway", "positive regulation of apoptotic process", "protein binding", "transmembrane transport" | Predicted to enable ATP:ADP antiporter activity. Involved in determination of adult lifespan; mitochondrion organization; and positive regulation of apoptotic process. Located in mitochondrion. Expressed in several structures, including body wall musculature; hermaphrodite somatic gonadal cell; intestine; pharynx; and vulva. Used to study mitochondrial metabolism disease. Human ortholog(s) of this gene implicated in Alzheimer's disease; autosomal dominant progressive external ophthalmoplegia with mitochondrial DNA deletions 2; facioscapulohumeral muscular dystrophy; intrinsic cardiomyopathy (multiple); and mitochondrial DNA depletion syndrome (multiple). |
| WBGene00007630 | *har-1* | C16C10.11 | CHCHD10 | "cytoplasm", "determination of adult lifespan", "mitochondrion organization", "mitochondrion", "nucleus", "striated muscle dense body" | Involved in mitochondrion organization. Acts upstream of with a positive effect on determination of adult lifespan. Located in mitochondrion and striated muscle dense body. Expressed in the intestine and ventral nerve cord. Human ortholog(s) of this gene implicated in isolated mitochondrial myopathy and neurodegenerative disease (multiple). |
| WBGene00008262 | *ril-1* | C53A5.1 | N.A. | "membrane" | Predicted to be located in the membrane. |
| WBGene00015248 | *mai-2* | B0546.1 | ATP5IF1 | "ATPase inhibitor activity", "cytoplasm", "mitochondrion", "negative regulation of ATP-dependent activity", "negative regulation of nucleotide metabolic process" | Enables ATPase inhibitor activity. Predicted to be involved in negative regulation of nucleotide metabolic process. Located in mitochondrion. Expressed in germ line and tail. |
| WBGene00017984 | *gmpr-1* | F32D1.5 | GMPR | "catalytic activity", "GMP reductase activity", "GMP reductase complex", "metal ion binding", "nucleotide metabolic process", "oxidoreductase activity", "purine nucleobase metabolic process", "purine nucleotide metabolic process" | Predicted to enable GMP reductase activity and metal ion binding activity. Predicted to be involved in purine nucleobase metabolic process and purine nucleotide metabolic process. Predicted to be part of GMP reductase complex. |

### 2.4 Experimental evidence supports the involvement of *ril-1* in OXPHOS

Among the five genes in the consensus list, *ril-1* emerged as the most compelling candidate for experimental validation because it lacks functional annotation beyond a predicted membrane localization. In addition, RNAi-mediated knockdown of *ril-1* has been reported to produce phenotypes commonly associated with impaired OXPHOS function, including larval arrest, embryonic lethality, slow growth, and reduced mitochondrial membrane potential.

To directly evaluate its potential role in mitochondrial respiration, we measured basal oxygen consumption in the *ril-1(ok2492)* heterozygous mutant strain. As shown in Figure 4, the mutant displayed reduced oxygen consumption relative to the N2 wild-type strain and values comparable to those observed in the *gas-1* heterozygous strain, which was included as a positive control for impaired mitochondrial respiration (39). These results provide experimental support for the involvement of RIL-1 in OXPHOS and independently validate the predictive power of the confidence-aware transcriptomic framework.

**Figure 4:**
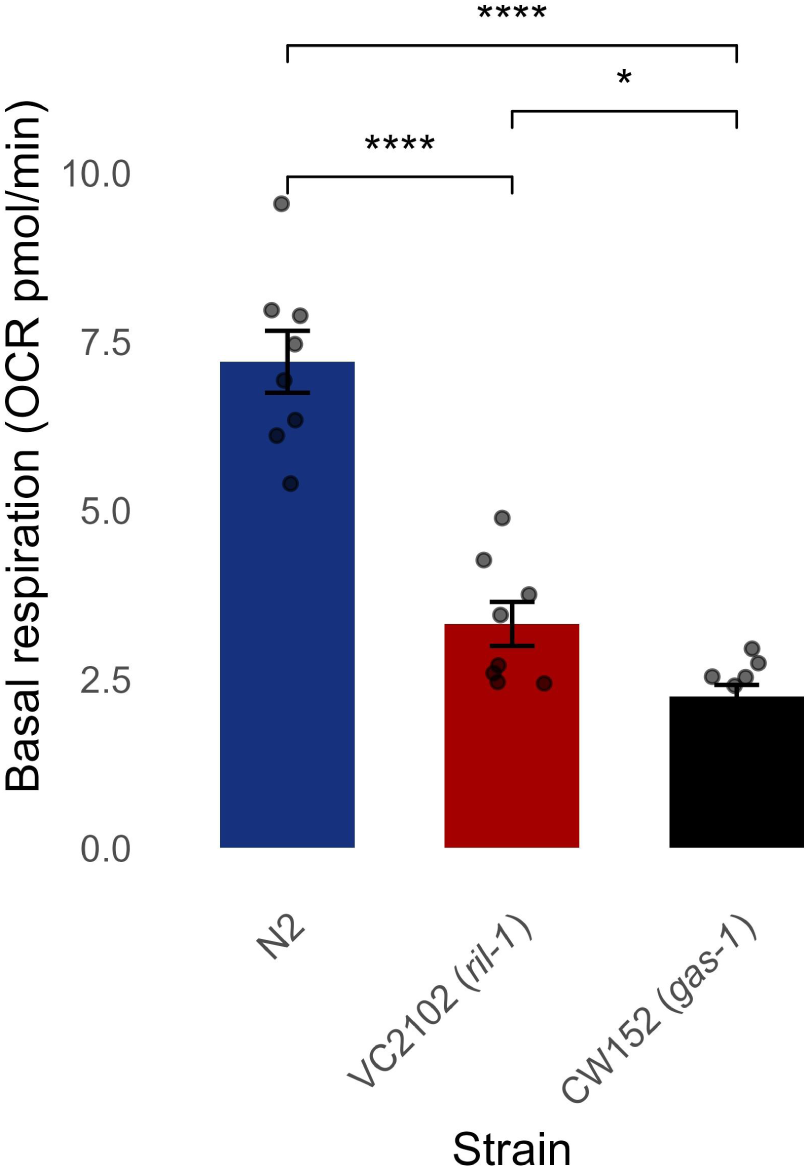
Reduced basal oxygen consumption in *ril-1* mutant animals. Basal oxygen consumption rate (OCR) was measured in wild-type (N2) animals, the heterozygous *ril-1(ok2492)* mutant strain VC2102, and the heterozygous *gas-1* mutant strain CW152, which served as a positive control for impaired mitochondrial respiration. The *ril-1* mutant exhibited reduced OCR relative to wild-type animals and values comparable to those of the *gas-1* control, supporting a defect in mitochondrial respiration. Asterisks indicate statistically significant differences (*P* < 0.05; \*\*\*\**P* < 0.0001) and bars represent mean ± SD. Data are representative of three independent biological replicates.

## 3 Discussion

Predicting gene function under incomplete and heterogeneous functional annotation remains a major challenge in functional genomics. In this study, we developed a confidence-aware learning framework that integrates supervised machine learning with transcriptome-based co-expression analysis to identify genes associated with oxidative phosphorylation (OXPHOS) in *C. elegans*. By combining multiple transcriptomic resolutions and requiring agreement across independent computational approaches, the framework prioritized a small set of high-confidence candidates while reducing the likelihood of predictions driven by biases specific to a single dataset or algorithm.

A notable result of the confidence-aware framework was that none of the genes annotated as respiratory complex assembly factors was recovered as a high-confidence candidate. This suggests that the transcriptional programs of assembly proteins differ from those of core and accessory OXPHOS components, supporting their exclusion from the initial positive training set and reinforcing the biological specificity of the learned transcriptional signature. These observations further suggest that assembly factors are not necessarily embedded within the same coordinated transcriptional module as mature OXPHOS components.

OXPHOS paralogs were intentionally excluded from the initial high-confidence OXPHOS set used in the first learning round because paralogous copies may not share the same transcriptional behavior as canonical OXPHOS genes. This design allowed us to evaluate whether the framework could independently recover the most OXPHOS-like member of each paralogous group. Consistent with this expectation, several paralogous families showed selective high-confidence recovery, with only one member exceeding the confidence threshold and being incorporated into the second learning round. Moreover, most of these recovered paralogs were also located within the OXPHOS-enriched co-expression modules identified from the single-cell transcriptomic analyses, providing independent support for their association with the OXPHOS transcriptional program.

A more nuanced pattern was observed for pairs in which none of them reached the probability threshold. In the case for complex I core subunit *gas-1*, this gene reached a probability of 0.72 while the paralog, *nduf-2.2*, reached a probability of 0.138. Similarly, the complex II core paralogs *sdha-1* and *sdha-2*, had probabilities of 0.588 and 0.116, respectively. For these cases, previous studies have indicated that the expression of these paralogous genes are inversely regulated, being *sdha-2* and *nduf-2.2* expressed almost exclusively in the germ line (40), while *gas-1* and *sdha-1* are expressed in most other cell types. These expression patterns can explain why neither paralog was recovered above the probability threshold, as well as the marked differences in predicted probabilities between the members of each paralogous pair. Similarly, paralogs such as *ucr-2.1, ucr-2.2* and *ucr-2.3* showed lower than threshold probabilities (0.777, 0.348 and 0.230, respectively), which may also be explained by a tissue-specific transcriptional regulation, as suggested by (41).

We also observed cases in which both paralogs showed probabilities close to the threshold. For example, *F45H10.2* and *R07E4.3* have probabilities of 0.897 and 0.847, respectively, suggesting that their expression profiles may partially resemble the OXPHOS-associated pattern but not strongly enough to satisfy the conservative criterion used here.

Together, these results suggest that most OXPHOS paralogs are not transcriptionally equivalent and that their recovery by the models depends on the specific regulatory context in which each copy is expressed. Therefore, although our approach was intentionally strict for training purposes, paralogs that did not reach the prediction threshold should not be interpreted as being unrelated to the OXPHOS system.

The conservative integration strategy adopted in this study substantially reduced the candidate space, yielding five genes supported simultaneously by the confidence-aware supervised framework and by OXPHOS-enriched co-expression modules in both developmental contexts. This consensus list provides a compact set of high-confidence candidates for experimental follow-up.

The final consensus list showed clear functional links to mitochondrial energy metabolism. *mai-2*, orthologous to human *ATP5IF1*, encodes ATPase inhibitory factor 1 (IF1), a key regulator of OXPHOS that prevents wasteful ATP hydrolysis under conditions of mitochondrial stress or low membrane potential (42,43). This gene is commonly considered an accessory regulator of the OXPHOS system (44). *ant-1.1*, orthologous to members of the mammalian *SLC25A4/5/6* family, encodes an adenine nucleotide translocator that couples mitochondrial ATP production to ADP/ATP exchange across the inner mitochondrial membrane (45,46). *har-1*, orthologous to human *CHCHD10*, is particularly relevant because CHCHD10 has been associated with mitochondrial cristae organization, respiratory chain integrity, and neuromuscular disease (47,48). Because respiratory complexes operate within the specialized architecture of the inner mitochondrial membrane, genes affecting mitochondrial structure may indirectly influence OXPHOS efficiency. In contrast, *gmpr-1*, a GMP reductase ortholog involved in purine nucleotide metabolism, represents a less direct but potentially informative candidate. Its recovery may reflect the coupling between respiration, nucleotide homeostasis, ATP demand, and cellular metabolic state (49,50). Together, these candidates indicate that the confidence-aware integration strategy can prioritize genes with both direct and indirect relationships to mitochondrial respiration, providing a focused set of hypotheses for future functional validation.

Among the consensus candidates, *ril-1* emerged as the strongest example of the type of discovery enabled by the confidence-aware framework. It remains poorly characterized and is currently annotated only as a membrane-associated protein, with no clear functional assignment. Importantly, RNAi-mediated knockdown of *ril-1* produces several phenotypes commonly associated with OXPHOS dysfunction, including larval arrest, embryonic lethality, slow growth, and reduced mitochondrial membrane potential. Together with its poor annotation and consistent prioritization across both supervised and co-expression analyses, these observations motivated its experimental characterization.

Because OXPHOS occurs in the inner mitochondrial membrane, the predicted membrane localization of RIL-1 is also compatible with a potential role in mitochondrial respiration. Unlike candidates supported by known orthology or established mitochondrial annotation, *ril-1* may represent genuinely novel biology that would be difficult to identify through homology-based approaches alone. Respirometry analysis of the *ril-1(ok2492)* heterozygous mutant revealed reduced oxygen consumption relative to the wild-type strain, providing direct experimental support for the involvement of RIL-1 in OXPHOS. Notably, *ril-1* appears to be restricted to the phylum Nematoda based on sequence and structural homology, being absent from mammals, *Drosophila*, and other animal lineages. Thus, *ril-1* emerges as a previously unrecognized nematode-specific OXPHOS candidate that would likely not have been identified through conventional homology-based annotation strategies.

Beyond the five consensus candidates, several additional genes from the supervised candidate set also merit further investigation. Among the top 10 genes in the supervised candidate set (Table S5), *F44E5.1*, *F23H11.5*, and *kdp-1* are also located within the adult co-expression module enriched in high-confidence OXPHOS genes. Notably, these genes currently lack experimentally validated functional annotations and display phenotypes associated with OXPHOS-related genes, while showing sequence homology only with genes from other nematodes. Thus, although their functional relationship with OXPHOS remains to be experimentally tested, their consistent recovery across independent computational approaches supports their prioritization as candidate genes potentially associated with OXPHOS or related mitochondrial processes.

Beyond the specific OXPHOS candidates identified here, our results demonstrate that transcriptomic landscapes can be systematically exploited for gene function prediction under incomplete functional annotation. The combination of confidence-aware supervised learning, informed bagging, and independent validation through single-cell co-expression analysis provides a principled strategy for extracting functional information from heterogeneous transcriptomic datasets. By explicitly distinguishing between high-confidence and weakly supported annotations, the framework incorporates biological uncertainty directly into the learning process, allowing candidate genes to be prioritized through convergent evidence rather than a single source of annotation. This design is especially relevant in functional genomics, where annotation quality is often heterogeneous.

This study also highlights the value of *C. elegans* as a discovery platform for mitochondrial biology. In addition to its rich genomic and transcriptomic resources, *C. elegans* offers powerful genetic tools that facilitate experimental validation of computational predictions. Furthermore, defining the complete repertoire of OXPHOS-associated genes has direct relevance for parasitic nematodes, which share several mitochondrial features with *C. elegans*, including an alternative electron transport chain that utilizes rhodoquinone instead of ubiquinone under hypoxic conditions (32–34). Because mitochondrial respiration has emerged as a promising antiparasitic target, identifying both conserved and lineage-specific OXPHOS regulators may have translational implications beyond the model organism. In this context, the recovery of the nematode-specific gene *ril-1* is particularly noteworthy, as it illustrates how transcriptome-based prediction can uncover lineage-restricted components that are largely inaccessible to homology-based annotation strategies.

In summary, we present a confidence-aware transcriptomic framework for functional gene discovery and apply it to oxidative phosphorylation in *C. elegans*. Our approach recovered genes with established bioenergetic roles while prioritizing poorly characterized candidates spanning transport, regulation, mitochondrial organization, and metabolic coupling. More broadly, this work illustrates how confidence-aware learning and multi-resolution transcriptomic integration can improve gene function prediction in biological systems where reliable functional annotations remain limited, providing a framework that may be transferable to other pathways, organisms, and functional genomics applications.

## 4 Methods

### 4.1 Overview of the computational framekork

We developed a confidence-aware learning framework that combines supervised machine learning with transcriptome-based co-expression analysis to predict genes associated with oxidative phosphorylation in *C. elegans*. The framework explicitly distinguishes between high-confidence OXPHOS genes, which are used for model training, and weakly supported candidates (low-confidence positives), which are incorporated only after independent computational support has been obtained.

The supervised component began with a curated set of high-confidence OXPHOS genes (Table S1). Twenty-five percent of these genes were withheld as an independent evaluation set, while the remaining genes were used to generate six alternative positive training sets: one containing genes from all OXPHOS complexes and five additional sets in which one respiratory complex was excluded at a time. This informed bagging strategy was designed to reduce potential biases associated with complex-specific transcriptional signatures.

For each positive training configuration, balanced negative sets were generated by randomly sampling genes from the remainder of the genome while excluding high-confidence OXPHOS genes (Table S1), low-confidence positives (Table S2), and respiratory complex assembly factors (Table S3). Three supervised learning algorithms (Random Forest, Support Vector Machine, and k-Nearest Neighbors) were trained on each positive/negative combination. Repeating the negative sampling procedure ten times produced a total of 180 binary classifiers.

Candidate genes were identified using a two-level ensemble voting strategy. Within each negative sampling iteration, a gene was considered positive if it was classified as positive by at least half of the 18 classifiers generated for that iteration. A second consensus step required a gene to be recovered in all ten independent negative sampling iterations, yielding a highly reproducible set of supervised candidates.

The low-confidence positives recovered with high confidence during the first learning round were subsequently incorporated into the positive training set for a second learning round, implementing the confidence-aware learning strategy illustrated in Figure 1. The resulting high-confidence supervised candidate set was then analyzed independently using single-cell transcriptomic data.

For the unsupervised component, embryonic and adult single-cell RNA-seq datasets were used to construct co-expression networks. Hierarchical clustering identified OXPHOS-enriched modules in each developmental context, and genes within these modules were considered unsupervised candidates according to the guilt-by-association principle (Table S6). Finally, the supervised candidate set was intersected with the OXPHOS-enriched modules from both embryonic and adult networks to generate the final consensus list of high-confidence OXPHOS candidates.

All computational analyses were performed using in-house Python and R code. Supervised learning models were implemented with the scikit-learn library unless otherwise stated.

### 4.2 Data

The computational framework integrated complementary transcriptomic resources representing two distinct levels of biological resolution. The supervised component was built using the developmental bulk RNA-seq atlas generated by Boeck et al. (51), which provides genome-wide expression profiles across multiple stages of the Caenorhabditis elegans life cycle, including a high-resolution embryonic time course and post-embryonic developmental stages.

The unsupervised component was built using single-cell RNA-seq datasets from Packer et al. (25) and Ghaddar et al. (52), representing embryonic and adult developmental contexts, respectively. For each dataset, gene expression values were averaged across cells assigned to the same annotated cell type, and the resulting cell-type-by-gene expression matrices were used for co-expression network construction and module enrichment analyses.

### 4.3 Sets of genes

#### High-confidence OXPHOS genes

A curated set of high-confidence OXPHOS genes was assembled from the literature and WormBase annotations. The final set comprised 64 genes encoding well-supported subunits of mitochondrial respiratory complexes I–V (Table S1). These genes constituted the initial positive training set for the supervised component of the framework.

To evaluate model generalization, 25% of the high-confidence OXPHOS genes were randomly withheld before training and used exclusively as an independent evaluation set in both rounds of training.

#### Low-confidence positives

Genes with evidence suggestive of OXPHOS function but insufficient to be included in the high-confidence training set were classified as low-confidence positives (Table S2). This category included two types of genes.

The first group consisted of 11 genes annotated in WormBase with GO terms related to oxidative phosphorylation based exclusively on indirect evidence codes (IEA or IBA). The second group comprised 25 duplicated or triplicated paralogs of canonical OXPHOS genes for which the functionally relevant copy could not be confidently determined from sequence information alone. These genes were excluded from the initial training set and were used to evaluate whether the framework could recover biologically plausible OXPHOS candidates during the confidence-aware learning procedure. Recovered genes with high confidence during the first learning round were incorporated into the positive training set for the second learning round

#### Assembly factor genes

Genes annotated as respiratory complex assembly factors were collected from WormBase and the literature (Table S3). These genes participate in the assembly or maturation of respiratory complexes but are not considered structural components of the mature OXPHOS machinery. They were excluded from both the high-confidence and low-confidence positive gene sets and from negative sampling during classifier construction.

#### Negative sampling pool

Negative examples were not defined as a fixed set of non-OXPHOS genes. Instead, for each training iteration, negative sets were generated by randomly sampling genes from the remainder of the *C. elegans* genome after excluding all high-confidence OXPHOS genes, low-confidence positives, and respiratory complex assembly factors. This strategy reduced the risk of introducing false negatives into the training process while allowing multiple alternative negative sets to be used during informed bagging.

### 4.4 Supervised learning Informed bagging strategy

The supervised component of the framework was designed to address two challenges: the limited number of high-confidence OXPHOS genes and the possibility that individual respiratory complexes exhibit transcriptional programs unrelated to canonical oxidative phosphorylation. Although complexes I–V function together in mitochondrial respiration, several complexes participate in additional processes, including redox homeostasis, reactive oxygen species signaling, hypoxia responses, apoptosis, and mitochondrial membrane organization. These complex-specific functions may introduce transcriptional signatures that are not exclusively associated with OXPHOS.

To reduce this potential source of bias, we implemented an informed bagging strategy inspired by bootstrap aggregation. Rather than generating bootstrap samples with replacement, we constructed six alternative positive training configurations from the high-confidence OXPHOS set: one containing genes from all respiratory complexes and five additional configurations in which one respiratory complex was excluded at a time (Figure 1). This strategy generated multiple partially independent views of the OXPHOS transcriptional program while preserving the biological structure of the pathway.

For each positive training configuration, balanced negative sets were generated by randomly sampling genes from the negative sampling pool defined in Section 4.3. The negative sampling procedure was repeated ten times, producing ten independent negative sets. Each positive configuration was combined with each negative set and used to train three supervised learning algorithms (Random Forest, Support Vector Machine, and k-Nearest Neighbors), resulting in a total of 180 binary classifiers (6 positive configurations × 10 negative sets × 3 algorithms).

#### Model training and evaluation

Supervised models were implemented in Python 3.10 using the pandas, numpy, matplotlib, and scikit-learn libraries. Hyperparameters were optimized using GridSearchCV with five-fold cross-validation and the F1-score as the optimization metric. The parameter grids explored for each algorithm are listed in Table S7.

Random Forest models were implemented using RandomForestClassifier(), Support Vector Machines using SVC(), and k-Nearest Neighbors using KNeighborsClassifier(). Because SVM and KNN are sensitive to feature scaling, gene expression values were standardized using StandardScaler() fitted on the training data and subsequently applied to the corresponding evaluation data.

Model performance was assessed using precision, recall, F1-score, and the area under the receiver operating characteristic curve (ROC AUC). Classification thresholds were optimized using precision-recall curves, selecting the probability threshold that maximized the F1-score on the evaluation set.

#### Ensemble voting and confidence-aware learning

Predictions from the 180 classifiers were integrated using a two-level ensemble voting strategy. Within each negative sampling iteration, a gene was considered positive if it was classified as positive by at least half of the 18 classifiers trained using that negative set (6 positive configurations × 3 algorithms). This produced one ensemble prediction for each of the ten independently sampled negative sets.

A second consensus step required a gene to be recovered in all ten ensemble predictions. Only genes satisfying this unanimous criterion were retained as high-confidence predictions from the first learning round. This conservative consensus strategy was designed to identify genes consistently associated with OXPHOS across multiple alternative positive configurations, negative sampling realizations, and learning algorithms.

The confidence-aware learning procedure consisted of two iterative learning rounds. The first round used the initial set of 48 high-confidence OXPHOS genes for training. Thirteen low-confidence positives recovered with a mean prediction probability greater than 0.9 were then incorporated into the positive training set, increasing its size from 48 to 61 genes. The complete informed bagging and ensemble procedure was repeated using this expanded training set.

The supervised candidate set used for downstream analyses was defined as the intersection of the predictions obtained in the first and second learning rounds, retaining only genes consistently recovered across both iterations.

### 4.5 Phenotype gene enrichment

Functional annotation coverage of the supervised candidate set was first assessed using Gene Ontology (GO) and KEGG annotations retrieved from Ensembl BioMart with the R package biomaRt. GO annotations were evaluated separately for the biological process, molecular function, and cellular component categories. Because a substantial fraction of the supervised candidates lacked informative GO or KEGG annotations, phenotype-based enrichment analysis was performed using WormBase phenotype annotations.

Phenotype annotations were obtained from the WormBase phenotype Gene Association File (GAF), release WBPS19. The file was parsed in R, retaining gene–phenotype associations with valid gene and phenotype identifiers and excluding annotations carrying the NOT qualifier. Duplicate gene–phenotype pairs were removed before downstream analyses.

Enrichment was evaluated using one-sided Fisher’s exact tests. The background was defined as all genes with at least one valid WormBase phenotype annotation, excluding the high-confidence OXPHOS genes used in the second learning round to avoid circularity. Fold enrichment was calculated as the ratio between the proportion of query genes annotated with a given phenotype and the corresponding proportion in the background, and p-values were adjusted using the Benjamini–Hochberg false discovery rate correction.

To define phenotype categories representative of canonical OXPHOS function, enrichment analysis was first performed using the high-confidence OXPHOS genes from the second learning round (61 genes). The ten most significantly enriched phenotype terms were selected as OXPHOS-associated phenotype categories and subsequently tested for enrichment in the supervised candidate set, considering genes recovered with a mean prediction probability greater than 0.9 in the second learning round.

### 4.6 Co-expression network analysis

To explore whether the predicted OXPHOS candidates were organized into transcriptionally coherent gene modules, we constructed two gene co-expression networks using single-cell RNA-seq-derived expression matrices from embryonic and adult *C. elegans*. For the embryonic network, we used the dataset from (25) summarizing expression by cell type independently of developmental time, resulting in 142 embryonic cell types. For the adult network, we used the dataset from (52), considering the annotated adult cell types, resulting in 180 cell types.

Co-expression networks were constructed in R using the WGCNA package following the general workflow described by (53). Briefly, genes and samples were filtered using the goodSamplesGenes function, soft-thresholding powers were selected according to the scale-free topology criterion, adjacency matrices were transformed into topological overlap matrices (TOMs), and genes were hierarchically clustered based on TOM dissimilarity. Gene modules were identified by dynamic tree cutting and summarized by module eigengenes. Modules with highly correlated eigengenes were merged using an eigengene dissimilarity threshold of 0.25.

For each network, we tested whether predefined OXPHOS-related gene sets were overrepresented within individual co-expression modules. The gene sets evaluated were the high-confidence OXPHOS genes used in the second learning round (61 genes), the low-confidence positives recovered after the first round of learning and the supervised candidate set recovered by the confidence-aware learning framework. Enrichment was assessed independently for each module using one-sided Fisher’s exact tests, with the genes present in the corresponding network used as background. Fold enrichment was calculated as the ratio between the proportion of genes from the query set within a module and the corresponding proportion in the network background. P-values were adjusted across modules using the Benjamini–Hochberg false discovery rate correction. For each module, we recorded the number of overlapping genes, fold enrichment, nominal p-value, adjusted FDR, and the identity of the overlapping genes.

### 4.7 Respirometry

Basal oxygen consumption rate (OCR) was measured in live *C. elegans* using a Seahorse XF Analyzer following the protocol described by Koopman et al. (39), with minor modifications. Age-synchronized young adult worms were collected in M9 buffer, washed twice to remove bacteria, and transferred to Seahorse assay plates containing a final volume of 200 µL of M9 buffer per well. Between 5 and 20 worms were loaded per well, and the exact number of animals in each well was determined after the assay to normalize OCR measurements.

Basal OCR was recorded using repeated cycles consisting of 2 min mixing, 2 min waiting, and 3 min measurement. OCR values were normalized to the number of worms in each well before statistical analysis.

## Author contributions

SZM; Conceptualization, Formal Analysis, Investigation, Methodology, Software, Review and Editing of the Manuscript. GS: Conceptualization, Investigation, Supervision, Review and Editing of the Manuscript. FPO: Conceptualization, Investigation, Methodology, Supervision, Writing of the Manuscript.

## Supplementary tables

Table S1 - High confidence OXPHOS genes

Table S2 - Low-confidence positives

Table S3 - Assembly factors

Table S4 - Predicted genes after 1st round of training

Table S5 - Supervised candidate set - Predicted genes by both rounds of learning

Table S6 - Genes in SC expression clusters highly enriched with high-confidence positives.xlsx

Table S7 - Explored parameters during supervised classifiers training

## Data availability statement

All code used is available on a GitHub repository at https://github.com/maszeballos/c-elegans-oxphos-gene-prediction.

We have also used Zenodo to assign a DOI to the repository: https://doi.org/10.5281/zenodo.21381998

The data used is available here: https://doi.org/10.5281/zenodo.21381998

