## Supplementary material for "Confidence-aware learning for transcriptome-based prediction of OXPHOS genes in *Caenorhabditis elegans* under incomplete functional annotation": Table S1

| Complex | gene_name_human | gene_name_worm | gene_ID_worm | Function | Evidence |
|---|---|---|---|---|---|
| I | MT-ND1 | nduo-1 | WBGene00010959 | Core | Tsang, W.Y., Lemire, B.D., 2003; Falk, M., 2009 |
| I | MT-ND2 | nduo-2 | WBGene00010961 | Core | Tsang, W.Y., Lemire, B.D., 2003; Falk, M., 2009 |
| I | MT-ND3 | nduo-3 | WBGene00010966 | Core | Tsang, W.Y., Lemire, B.D., 2003; Falk, M., 2009 |
| I | MT-ND4 | nduo-4 | WBGene00010963 | Core | Tsang, W.Y., Lemire, B.D., 2003; Falk, M., 2009 |
| I | MT-ND4L | ndfl-4 | WBGene00010958 | Core | Tsang, W.Y., Lemire, B.D., 2003; Falk, M., 2009 |
| I | MT-ND5 | nduo-5 | WBGene00010967 | Core | Tsang, W.Y., Lemire, B.D., 2003; Falk, M., 2009 |
| I | MT-ND6 | nduo-6 | WBGene00010957 | Core | Tsang, W.Y., Lemire, B.D., 2003; Falk, M., 2009 |
| I | NDUFA10 | nuo-4 | WBGene00019401 | Accesory | Tsang, W.Y., Lemire, B.D., 2003; Falk, M., 2009 |
| I | NDUFA11 | nduf-11 | WBGene00007192 | Accesory | Knapp-Wilson, A 2021 |
| I | NDUFA12 | ndua-12 | WBGene00022380 | Accesory | Tsang, W.Y., Lemire, B.D., 2003; Falk, M., 2009 |
| I | NDUFA13 | ndua-13 | WBGene00016393 | Accesory | Tsang, W.Y., Lemire, B.D., 2003; Falk, M., 2009 |
| I | NDUFA2 | ndua-2 | WBGene00013406 | Core | Tsang, W.Y., Lemire, B.D., 2003; Falk, M., 2009 |
| I | NDUFA5 | ndua-5 | WBGene00007880 | Accesory | Tsang, W.Y., Lemire, B.D., 2003; Falk, M., 2009 |
| I | NDUFA6 | nuo-3 | WBGene00013308 | Core | Tsang, W.Y., Lemire, B.D., 2003; Falk, M., 2009 |
| I | NDUFA7 | ndua-7 | WBGene00009740 | Accesory | Tsang, W.Y., Lemire, B.D., 2003; Falk, M., 2009 |
| I | NDUFA8 | ndua-8 | WBGene00021849 | Accesory | Tsang, W.Y., Lemire, B.D., 2003; Falk, M., 2009 |
| I | NDUFA9 | nduf-9 | WBGene00021800 | Accesory | Tsang, W.Y., Lemire, B.D., 2003; Falk, M., 2009 |
| I | NDUFB10 | ndub-10 | WBGene00010326 | Accesory | Tsang, W.Y., Lemire, B.D., 2003; Falk, M., 2009 |
| I | NDUFB11 | ndub-11 | WBGene00018361 | Accesory | Falk, M., 2009 |
| I | NDUFB2 | ndub-2 | WBGene00009712 | Accesory | Tsang, W.Y., Lemire, B.D., 2003; Falk, M., 2009 |
| I | NDUFB4 | nuo-6 | WBGene00012166 | Core | Tsang, W.Y., Lemire, B.D., 2003; Falk, M., 2009 |
| I | NDUFB5 | ndub-5 | WBGene00016118 | Accesory | Tsang, W.Y., Lemire, B.D., 2003; Falk, M., 2009 |
| I | NDUFB6 | ndub-6 | WBGene00014086 | Core | Falk, M., 2009 |
| I | NDUFB7 | ndub-7 | WBGene00008414 | Accesory | Tsang, W.Y., Lemire, B.D., 2003; Falk, M., 2009 |
| I | NDUFB8 | ndub-8 | WBGene00013094 | Accesory | Tsang, W.Y., Lemire, B.D., 2003; Falk, M., 2009 |
| I | NDUFB9 | ndub-9 | WBGene00015810 | Accesory | Tsang, W.Y., Lemire, B.D., 2003; Falk, M., 2009 |
| I | NDUFC2 | nduc-2 | WBGene00022169 | Accesory | Tsang, W.Y., Lemire, B.D., 2003; Falk, M., 2009 |
| I | NDUFS1 | nuo-5 | WBGene00021562 | Core | Tsang, W.Y., Lemire, B.D., 2003; Falk, M., 2009 |
| I | NDUFS3 | nuo-2 | WBGene00020417 | Core | Tsang, W.Y., Lemire, B.D., 2003; Falk, M., 2009 |

| | | | | | |
|---|---|---|---|---|---|
| I | NDUFS4 | lpd-5 | WBGene00003061 | Accesory | Tsang, W.Y., Lemire, B.D., 2003; Falk, M., 2009 |
| I | NDUFS5 | nduf-5 | WBGene00021839 | Accesory | Tsang, W.Y., Lemire, B.D., 2003; Falk, M., 2009 |
| I | NDUFS6 | nduf-6 | WBGene00009051 | Accesory | Tsang, W.Y., Lemire, B.D., 2003; Falk, M., 2009 |
| I | NDUFS7 | nduf-7 | WBGene00012376 | Core | Tsang, W.Y., Lemire, B.D., 2003; Falk, M., 2009 |
| I | NDUFS8 | ndus-8 | WBGene00020636 | Core | Tsang, W.Y., Lemire, B.D., 2003; Falk, M., 2009 |
| I | NDUFV1 | nuo-1 | WBGene00003831 | Core | Tsang, W.Y., Lemire, B.D., 2003; Falk, M., 2009 |
| I | NDUFV2 | nduv-2 | WBGene00009992 | Core | Tsang, W.Y., Lemire, B.D., 2003; Falk, M., 2009 |
| II | SDHB | sdhb-1 | WBGene00006433 | Core | Tsang, W.Y., Lemire, B.D., 2003 |
| II | SDHC | mev-1 | WBGene00003225 | Core | Tsang, W.Y., Lemire, B.D., 2003 |
| II | SDHD | sdhd-1 | WBGene00009353 | Core | Tsang, W.Y., Lemire, B.D., 2003 |
| III | CYC1 | cyc-1 | WBGene00000869 | Core | Tsang, W.Y., Lemire, B.D., 2003 |
| III | MT-CYB | ctb-1 | WBGene00000829 | Core | Tsang, W.Y., Lemire, B.D., 2003 |
| III | UQCRB | T02H6.11 | WBGene00020181 | Core | Tsang, W.Y., Lemire, B.D., 2003 |
| III | UQCRFS1 | isp-1 | WBGene00002162 | Core | Tsang, W.Y., Lemire, B.D., 2003 |
| III | UQCRH | T27E9.2 | WBGene00012094 | Core | Tsang, W.Y., Lemire, B.D., 2003 |
| IV | COX4I1 | cox-4 | WBGene00012354 | Core | Tsang, W.Y., Lemire, B.D., 2003 |
| IV | COX5A | cox-5a | WBGene00012553 | Core | Tsang, W.Y., Lemire, B.D., 2003 |
| IV | COX5B | cox-5b | WBGene00000371 | Core | Tsang, W.Y., Lemire, B.D., 2003 |
| IV | COX6A1, COX6A2 | cox-6a | WBGene00006519 | Core | Tsang, W.Y., Lemire, B.D., 2003 |
| IV | COX6B1 | cox-6b | WBGene00022170 | Core | Tsang, W.Y., Lemire, B.D., 2003 |
| IV | COX7C | cox-7c | WBGene00009161 | Core | Tsang, W.Y., Lemire, B.D., 2003 |
| IV | MT-CO1 | ctc-1 | WBGene00010964 | Core | Tsang, W.Y., Lemire, B.D., 2003 |
| IV | MT-CO2 | ctc-2 | WBGene00010965 | Core | Tsang, W.Y., Lemire, B.D., 2003 |
| IV | MT-CO3 | ctc-3 | WBGene00010962 | Core | Tsang, W.Y., Lemire, B.D., 2003 |
| V | ATP5F1A | atp-1 | WBGene00010419 | Core | Tsang, W.Y., Lemire, B.D., 2003 |
| V | ATP5F1B | atp-2 | WBGene00000229 | Core | Tsang, W.Y., Lemire, B.D., 2003 |
| V | ATP5F1C | Y69A2AR.18 | WBGene00022089 | Core | Tsang, W.Y., Lemire, B.D., 2003 |
| V | ATP5F1D | F58F12.1 | WBGene00019061 | Core | Tsang, W.Y., Lemire, B.D., 2003 |
| V | ATP5MC1, ATP5MC [illegible] | Y82E9BR.3 | WBGene00022336 | Core | Tsang, W.Y., Lemire, B.D., 2003 |
| V | ATP5ME | R04F11.2 | WBGene00011015 | Core | Tsang, W.Y., Lemire, B.D., 2003 |

| | | | | | |
|---|---|---|---|---|---|
| V | ATP5MF | R53.4 | WBGene00011273 | Core | Tsang, W.Y., Lemire, B.D., 2003 |
| V | ATP5PD | atp-5 | WBGene00007385 | Core | Tsang, W.Y., Lemire, B.D., 2003 |
| V | ATP5PF | atp-4 | WBGene00020275 | Core | Human otholog |
| V | ATP5PO | atp-3 | WBGene00000230 | Core | Tsang, W.Y., Lemire, B.D., 2003 |
| V | ATPAF2 | Y116A8C.27 | WBGene00013804 | Core | Human otholog |
| V | MT-ATP6 | atp-6 | WBGene00010960 | Core | Tsang, W.Y., Lemire, B.D., 2003 |
