## Supplementary material for "Confidence-aware learning for transcriptome-based prediction of OXPHOS genes in *Caenorhabditis elegans* under incomplete functional annotation": Table S2

| Complex | Gene name | WormBase ID | Human Orthologous | GO Annotation | 1st round with mean probability > 0.9 |
|---|---|---|---|---|---|
| **Weak evidence** | | | | | |
| I | *nduv-3* | WBGene00022075 | NDUFV3 | *GO:0045271; IEA* | NO |
| I | *C06A5.3* | WBGene00015501 | HDGF | *GO:0045271; IEA* | NO |
| I | *ndub-3* | WBGene00007684 | NDUFB3 | *GO:0045271; IBA, IEA* | YES |
| I | ndua-1 | WBGene00009294 | NDUFA1 | *GO:0045271; ISS* | YES |
| III | *ucr-11* | WBGene00019007 | UQCR11 | *GO:0006122; IEA* | YES |
| III | *C14B9.10* | WBGene00015755 | UQCR10 | *GO:0006122; IEA* | YES |
| IV | F36A2.7 | WBGene00009454 | NDUA4 | NA | YES |
| IV | F29B9.11 | WBGene00017925 | NA | NA | YES |
| **Paralogues** | | | | | |
| I | *ndab-1* | WBGene00013237 | NDUFAB1 | | NO |
| I | *ndab-2* | WBGene00018151 | NDUFAB1 | | YES |
| I | *gas-1* | WBGene00001520 | NDUFS2 | | NO |
| I | *nduf2-2* | WBGene00006463 | NDUFS2 | | NO |
| II | *sdha-1* | WBGene00015391 | SDHA | | NO |
| II | *sdha-2* | WBGene00016392 | SDHA | | NO |
| III | *F45H10.2* | WBGene00009739 | UQCRQ | | NO |
| III | *R07E4.3* | R07E4.3 | UQCRQ | | NO |
| III | *mppb-1* | WBGene00013880 | UQCRC1 | | NO |
| III | *ucr-1* | WBGene00018963 | UQCRC1 | | YES |
| III | *ucr-2.1* | WBGene00012158 | UQCRC2 | | NO |
| III | *ucr-2.2* | WBGene00011679 | UQCRC2 | | NO |
| III | *ucr-2.3* | ucr-2.3 | UQCRC2 | | NO |
| IV | *Y111B2A.2* | WBGene00013728 | COX6C | | NO |
| IV | *B0035.18* | B0035.18 | COX6C | | NO |
| IV | *cox-6c* | WBGene00017926 | COX6C | | YES |
| IV | *cyc-2.1* | WBGene00017121 | CYS1 | | YES |
| IV | *cyc-2.2* | WBGene00013854 | CYS1 | | NO |
| V | *asb-1* | WBGene00000206 | ATP5PB | | NO |
| V | *asb-2* | WBGene00000207 | ATP5PB | | YES |
| V | *asg-1* | WBGene00000209 | ATP5MG | | NO |
| V | *asg-2* | WBGene00000210 | ATP5MG | | YES |
| V | *hpo-18* | WBGene00017982 | ATP5F1E | | YES |
| V | *R05D3.6* | WBGene00019880 | ATP5F1E | | NO |
| V | *ZC262.5* | WBGene00022582 | ATP5F1E | | NO |
