## Supplementary material for "Confidence-aware learning for transcriptome-based prediction of OXPHOS genes in *Caenorhabditis elegans* under incomplete functional annotation": Table S3

| Complex | Gene name | WBID | Human Orthologue |
|---|---|---|---|
| I | acdh-12 | WBGene00012860 | ACADVL |
| I | B0035.15 | WBGene00007113 | NDUFAF4 |
| I | B0334.5 | WBGene00007145 | NDUFAF6 |
| I | K09E4.3 | WBGene00010721 | NDUFAF5 |
| I | M04B2.4 | WBGene00010847 | FOXRD1 |
| I | ndua-2 | WBGene00013406 | NDUFA2 |
| I | nuaf-1 | WBGene00008225 | NDUFAF1 |
| I | nuaf-3 | WBGene00011123 | NDUFAF3 |
| I | nubp-1 | WBGene00008664 | NUBP1 |
| I | Y116A8C.30 | WBGene00013807 | NDUFAF2 |
| I | Y38F2AR.3 | WBGene00021421 | TIMMDC1 |
| I | ZK1128.1 | WBGene00014227 | NDUFAF7 |
| II | Y57A10A.29 | WBGene00013269 | SDHAF2 |
| III | bcs-1 | WBGene00010042 | BCS1L |
| III | ddl-1 | WBGene00019125 | TTC19 |
| IV | coa-1 | WBGene00195063 | COA1 |
| IV | coa-3 | WBGene00018046 | COA3 |
| IV | coa-4 | WBGene00011857 | COA4 |
| IV | coa-5 | WBGene00012483 | COA5 |
| IV | coa-6 | WBGene00013893 | PRORP |
| IV | coa-7 | WBGene00013925 | COA7 |
| IV | cox-10 | WBGene00012895 | COX10 |
| IV | cox-11 | WBGene00010437 | COX11 |
| IV | cox-14 | WBGene00011875 | COX14 |
| IV | cox-15 | WBGene00011526 | COX15 |
| IV | cox-16 | WBGene00302991 | COX16 |
| IV | cox-17 | WBGene00018240 | COX17 |
| IV | cox-18 | WBGene00021932 | COX18 |
| IV | cox-19 | WBGene00009745 | COX19 |
| IV | sco-1 | WBGene00015297 | SCO1 |
| IV | stf-1 | WBGene00004787 | SURF1 |
| IV | T20D3.6 | WBGene00011859 | HIGD2A |
| IV | Y53F4B.14 | WBGene00013161 | PET100 |
