## Supplementary material for "Confidence-aware learning for transcriptome-based prediction of OXPHOS genes in *Caenorhabditis elegans* under incomplete functional annotation": Table S5

| Worm Base ID | Gene name | Avg. Probability (both iterations) |
|---|---|---|
| WBGene00009688 | F44E5.1 | 0,97 |
| WBGene00006911 | vha-2 | 0,96 |
| WBGene00008262 | ril-1 | 0,96 |
| WBGene00000263 | F23H11.5 | 0,96 |
| WBGene00019322 | ahcy-1 | 0,96 |
| WBGene00013463 | kdp-1 | 0,96 |
| WBGene00006434 | prdx-2 | 0,96 |
| WBGene00000065 | act-3 | 0,96 |
| WBGene00000063 | act-1 | 0,96 |
| WBGene00000833 | cts-1 | 0,96 |
| WBGene00002344 | let-70 | 0,96 |
| WBGene00003920 | par-5 | 0,96 |
| WBGene00000883 | cyn-7 | 0,96 |
| WBGene00019900 | vdac-1 | 0,96 |
| WBGene00002045 | icd-1 | 0,96 |
| WBGene00000802 | crt-1 | 0,96 |
| WBGene00008505 | F01G4.6 | 0,96 |
| WBGene00001005 | dlc-1 | 0,96 |
| WBGene00001427 | fkb-2 | 0,95 |
| WBGene00006537 | tbb-2 | 0,95 |
| WBGene00009119 | ndk-1 | 0,95 |
| WBGene00000064 | act-2 | 0,95 |
| WBGene00018519 | F46H5.3 | 0,95 |
| WBGene00017166 | aldo-2 | 0,95 |
| WBGene00011156 | rbm-3.2 | 0,95 |
| WBGene00006528 | tba-1 | 0,95 |
| WBGene00008920 | eef-1G | 0,95 |
| WBGene00018846 | eef-1B.1 | 0,95 |
| WBGene00006536 | tbb-1 | 0,95 |
| WBGene00019537 | K08D12.3 | 0,95 |
| WBGene00004435 | rpl-23 | 0,95 |
| WBGene00006924 | vig-1 | 0,94 |
| WBGene00006919 | vha-10 | 0,94 |
| WBGene00001502 | ftt-2 | 0,94 |
| WBGene00006529 | tba-2 | 0,94 |
| WBGene00006439 | ant-1.1 | 0,94 |
| WBGene00009880 | F49C12.11 | 0,94 |
| WBGene00003964 | pdi-3 | 0,94 |
| WBGene00015248 | mai-2 | 0,94 |
| WBGene00022235 | sqd-1 | 0,94 |
| WBGene00002980 | lgg-1 | 0,94 |
| WBGene00016746 | C48B6.10 | 0,94 |
| WBGene00022748 | oaz-1 | 0,94 |
| WBGene00000380 | cct-5 | 0,94 |

| | | |
|---|---|---|
| WBGene00003902 | pab-1 | 0,94 |
| WBGene00009122 | tct-1 | 0,93 |
| WBGene00004424 | rpl-12 | 0,93 |
| WBGene00020868 | eif-1 | 0,93 |
| WBGene00016655 | acbp-1 | 0,93 |
| WBGene00004471 | rps-2 | 0,93 |
| WBGene00010809 | lias-1 | 0,93 |
| WBGene00000475 | cey-4 | 0,93 |
| WBGene00001168 | eef-1A.1 | 0,93 |
| WBGene00011128 | adk-1 | 0,93 |
| WBGene00011232 | pck-2 | 0,93 |
| WBGene00006917 | vha-8 | 0,93 |
| WBGene00012602 | Y38E10A.24 | 0,93 |
| WBGene00001303 | sec-61.G | 0,93 |
| WBGene00009050 | mstr-1 | 0,93 |
| WBGene00001971 | hmg-1.1 | 0,93 |
| WBGene00020894 | T28D9.1 | 0,93 |
| WBGene00077526 | C25A1.16 | 0,93 |
| WBGene00004474 | rps-5 | 0,93 |
| WBGene00007708 | nola-3 | 0,93 |
| WBGene00006910 | vha-1 | 0,93 |
| WBGene00002005 | hsp-1 | 0,93 |
| WBGene00000881 | cyn-5 | 0,93 |
| WBGene00004442 | rpl-28 | 0,92 |
| WBGene00004450 | rpl-36 | 0,92 |
| WBGene00001946 | his-72 | 0,92 |
| WBGene00004439 | rpl-23A.2 | 0,92 |
| WBGene00000879 | cyn-3 | 0,92 |
| WBGene00012179 | rpl-37.1 | 0,92 |
| WBGene00000066 | act-4 | 0,92 |
| WBGene00004484 | rps-15 | 0,92 |
| WBGene00004888 | smo-1 | 0,92 |
| WBGene00004489 | rps-20 | 0,92 |
| WBGene00004434 | rpl-22 | 0,92 |
| WBGene00004302 | ran-1 | 0,92 |
| WBGene00004456 | rpl-37A | 0,92 |
| WBGene00004479 | rps-10 | 0,92 |
| WBGene00006984 | zig-7 | 0,92 |
| WBGene00004488 | rps-19 | 0,92 |
| WBGene00004492 | rps-23 | 0,92 |
| WBGene00004493 | rps-24 | 0,92 |
| WBGene00003065 | lpd-9 | 0,92 |
| WBGene00017075 | nap-1 | 0,92 |
| WBGene00003934 | pat-10 | 0,92 |
| WBGene00021248 | Y22D7AL.10 | 0,92 |
| WBGene00010896 | snu-13 | 0,92 |

| | | |
|---|---|---|
| WBGene00004451 | rpl-37.2 | 0,92 |
| WBGene00004410 | rplp-2.2 | 0,92 |
| WBGene00009882 | vha-17 | 0,92 |
| WBGene00000183 | arf-5 | 0,92 |
| WBGene00019680 | K12H4.5 | 0,92 |
| WBGene00001911 | his-37 | 0,92 |
| WBGene00004478 | rps-9 | 0,92 |
| WBGene00017984 | gmpr-1 | 0,92 |
| WBGene00004897 | snb-1 | 0,92 |
| WBGene00003053 | lmp-1 | 0,92 |
| WBGene00004428 | rpl-13A | 0,92 |
| WBGene00003962 | pdi-1 | 0,91 |
| WBGene00001169 | eef-1A.2 | 0,91 |
| WBGene00007630 | har-1 | 0,91 |
| WBGene00006728 | ubq-2 | 0,91 |
| WBGene00004447 | rpl-35A | 0,91 |
| WBGene00014016 | ZK632.9 | 0,91 |
| WBGene00012344 | ola-1 | 0,91 |
| WBGene00016258 | vha-16 | 0,91 |
| WBGene00006725 | rps-27A | 0,91 |
| WBGene00004472 | rps-3 | 0,91 |
| WBGene00004408 | rplp-0 | 0,91 |
| WBGene00004419 | rpl-7A | 0,91 |
| WBGene00019162 | eif-1.A | 0,91 |
| WBGene00004426 | rpl-14 | 0,91 |
| WBGene00006727 | ubq-1 | 0,91 |
| WBGene00016493 | rplp-2.1 | 0,91 |
| WBGene00021350 | rpl-27A | 0,91 |
| WBGene00000377 | cct-1 | 0,91 |
| WBGene00004487 | rps-18 | 0,91 |
| WBGene00004436 | rpl-24 | 0,91 |
| WBGene00001558 | gdi-1 | 0,91 |
| WBGene00010478 | dkc-1 | 0,91 |
| WBGene00004444 | rpl-30 | 0,91 |
| WBGene00004441 | rpl-27 | 0,91 |
| WBGene00001748 | gsp-2 | 0,91 |
| WBGene00004429 | rpl-17 | 0,91 |
| WBGene00006913 | vha-4 | 0,91 |
| WBGene00004415 | rpl-4 | 0,91 |
| WBGene00013766 | prmt-1 | 0,91 |
| WBGene00004480 | rps-11 | 0,91 |
| WBGene00004483 | rps-14 | 0,91 |
| WBGene00004421 | rpl-10L | 0,91 |
| WBGene00004409 | rplp-1 | 0,91 |
| WBGene00004431 | rpl-19 | 0,91 |
| WBGene00001167 | eef-2 | 0,91 |

| | | |
|---|---|---|
| WBGene00020604 | T20B12.7 | 0,91 |
| WBGene00003963 | pdi-2 | 0,91 |
| WBGene00004469 | rpsa-1 | 0,91 |
| WBGene00009772 | ztf-7 | 0,91 |
| WBGene00001423 | fib-1 | 0,91 |
| WBGene00003372 | mlc-4 | 0,90 |
| WBGene00006918 | vha-9 | 0,90 |
| WBGene00004920 | snr-7 | 0,90 |
| WBGene00004494 | rps-25 | 0,90 |
| WBGene00004423 | rpl-11.2 | 0,90 |
| WBGene00022599 | daf-41 | 0,90 |
| WBGene00004449 | rpl-35 | 0,90 |
| WBGene00001840 | hel-1 | 0,90 |
| WBGene00015514 | srlf-1 | 0,90 |
| WBGene00001501 | ftn-2 | 0,90 |
| WBGene00004418 | rpl-7 | 0,90 |
| WBGene00004470 | rps-3A | 0,90 |
| WBGene00019831 | R02F2.1 | 0,90 |
| WBGene00014095 | gdh-1 | 0,90 |
| WBGene00004432 | rpl-18A | 0,90 |
| WBGene00004496 | rps-27 | 0,90 |
| WBGene00004476 | rps-7 | 0,90 |
| WBGene00004445 | rpl-31 | 0,90 |
| WBGene00000896 | dad-1 | 0,90 |
| WBGene00004477 | rps-8 | 0,90 |
| WBGene00004412 | rpl-10a | 0,90 |
| WBGene00004446 | rpl-32 | 0,90 |
| WBGene00022783 | tomm-7 | 0,90 |
| WBGene00004433 | rpl-21 | 0,90 |
| WBGene00004440 | rpl-26 | 0,90 |
| WBGene00004482 | rps-13 | 0,90 |
| WBGene00004143 | pqn-59 | 0,90 |
| WBGene00004417 | rpl-6 | 0,90 |
| WBGene00004430 | rpl-18 | 0,90 |
| WBGene00004499 | fau-1 | 0,90 |
| WBGene00002007 | hsp-3 | 0,90 |
| WBGene00004700 | rsp-3 | 0,90 |
| WBGene00002191 | kin-3 | 0,89 |
| WBGene00000182 | arf-1 | 0,89 |
| WBGene00004914 | snr-1 | 0,89 |
| WBGene00010556 | rack-1 | 0,89 |
| WBGene00004420 | rpl-9 | 0,89 |
| WBGene00018491 | mdh-1 | 0,89 |
| WBGene00004498 | rps-29 | 0,89 |
| WBGene00004416 | rpl-5 | 0,89 |
| WBGene00004481 | rps-12 | 0,89 |

| | | |
|---|---|---|
| WBGene00004414 | rpl-3 | 0,89 |
| WBGene00004486 | rps-17 | 0,89 |
| WBGene00000534 | cpi-2 | 0,89 |
| WBGene00022127 | yop-1 | 0,89 |
| WBGene00004452 | rpl-38 | 0,89 |
| WBGene00004491 | rps-15A | 0,89 |
| WBGene00004438 | rpl-23A.1 | 0,89 |
| WBGene00004425 | rpl-13 | 0,89 |
| WBGene00004918 | snr-5 | 0,89 |
| WBGene00017675 | F21F3.6 | 0,89 |
| WBGene00004454 | rpl-36A | 0,89 |
| WBGene00004448 | rpl-34 | 0,89 |
| WBGene00004473 | rps-4 | 0,89 |
| WBGene00013238 | trap-4 | 0,89 |
| WBGene00004475 | rps-6 | 0,89 |
| WBGene00004490 | rps-21 | 0,89 |
| WBGene00006730 | uev-1 | 0,89 |
| WBGene00000915 | hsp-90 | 0,89 |
| WBGene00022042 | icd-2 | 0,89 |
| WBGene00003370 | mlc-2 | 0,89 |
| WBGene00010627 | nol-56 | 0,89 |
| WBGene00001685 | gpd-3 | 0,89 |
| WBGene00001749 | gst-1 | 0,89 |
| WBGene00004766 | sel-9 | 0,88 |
| WBGene00003123 | mag-1 | 0,88 |
| WBGene00003901 | paa-1 | 0,88 |
| WBGene00004427 | rpl-15 | 0,88 |
| WBGene00012097 | abcf-2 | 0,88 |
| WBGene00004046 | plp-1 | 0,88 |
| WBGene00007235 | ahsa-1 | 0,88 |
| WBGene00000472 | cey-1 | 0,88 |
| WBGene00018784 | F54A3.5 | 0,88 |
| WBGene00020683 | ribo-1 | 0,88 |
| WBGene00010130 | vha-14 | 0,88 |
| WBGene00006574 | tin-13 | 0,88 |
| WBGene00002083 | inf-1 | 0,88 |
| WBGene00012351 | W09C5.1 | 0,88 |
| WBGene00007223 | znf-706 | 0,88 |
| WBGene00018395 | mtch-1 | 0,88 |
| WBGene00010337 | serp-1.1 | 0,88 |
| WBGene00002202 | kin-19 | 0,88 |
| WBGene00004485 | rps-16 | 0,88 |
| WBGene00013311 | sec-61.A | 0,88 |
| WBGene00004453 | rpl-39 | 0,88 |
| WBGene00004495 | rps-26 | 0,88 |
| WBGene00009881 | F49C12.12 | 0,88 |

| | | |
|---|---|---|
| WBGene00002065 | iff-2 | 0,87 |
| WBGene00004443 | rpl-29 | 0,87 |
| WBGene00015168 | pdi-6 | 0,87 |
| WBGene00020216 | trap-2 | 0,87 |
| WBGene00002196 | kin-10 | 0,87 |
| WBGene00000776 | cpl-1 | 0,87 |
| WBGene00000198 | art-1 | 0,87 |
| WBGene00004497 | rps-28 | 0,87 |
| WBGene00000941 | ddp-1 | 0,87 |
| WBGene00000552 | cmd-1 | 0,87 |
| WBGene00006704 | ubc-7 | 0,87 |
| WBGene00003214 | mel-32 | 0,87 |
| WBGene00006959 | xbp-1 | 0,87 |
| WBGene00004930 | sod-1 | 0,87 |
| WBGene00004422 | rpl-11.1 | 0,87 |
| WBGene00021960 | tmem-258 | 0,87 |
| WBGene00006601 | tpi-1 | 0,87 |
| WBGene00004413 | rpl-8 | 0,87 |
| WBGene00004274 | rab-11.1 | 0,86 |
| WBGene00010333 | copz-1 | 0,86 |
| WBGene00011110 | prdx-3 | 0,86 |
| WBGene00001137 | eat-6 | 0,86 |
| WBGene00010260 | ddx-17 | 0,86 |
| WBGene00003989 | pfn-1 | 0,86 |
| WBGene00000779 | cpn-3 | 0,86 |
| WBGene00020931 | cytb-5.2 | 0,86 |
| WBGene00001037 | dnj-19 | 0,86 |
| WBGene00022053 | cisd-3.2 | 0,86 |
| WBGene00021088 | W08E12.7 | 0,86 |
| WBGene00004917 | snr-4 | 0,86 |
| WBGene00001684 | gpd-2 | 0,86 |
| WBGene00004919 | snr-6 | 0,86 |
| WBGene00011884 | enol-1 | 0,86 |
| WBGene00000067 | act-5 | 0,86 |
| WBGene00006921 | vha-12 | 0,86 |
| WBGene00000235 | baf-1 | 0,86 |
| WBGene00020112 | pfd-5 | 0,86 |
| WBGene00003893 | ost-1 | 0,86 |
| WBGene00020297 | nog-1 | 0,86 |
| WBGene00019466 | tos-1 | 0,85 |
| WBGene00006833 | unc-108 | 0,85 |
| WBGene00009664 | idha-1 | 0,85 |
| WBGene00001385 | far-1 | 0,85 |
| WBGene00002010 | hsp-6 | 0,85 |
| WBGene00006713 | ubc-18 | 0,85 |
| WBGene00020915 | nol-58 | 0,85 |

| | | |
|---|---|---|
| WBGene00001564 | icl-1 | 0,85 |
| WBGene00077771 | tspo-1 | 0,85 |
| WBGene00012964 | Y48A6B.3 | 0,85 |
| WBGene00002264 | lec-1 | 0,85 |
| WBGene00000114 | alh-8 | 0,85 |
| WBGene00021427 | sec-61.B | 0,85 |
| WBGene00002978 | lev-11 | 0,85 |
| WBGene00004916 | snr-3 | 0,85 |
| WBGene00003371 | mlc-3 | 0,85 |
| WBGene00003369 | mlc-1 | 0,84 |
| WBGene00010303 | cri-3 | 0,84 |
| WBGene00021024 | W04C9.2 | 0,84 |
| WBGene00011480 | enpl-1 | 0,84 |
| WBGene00004015 | phb-2 | 0,84 |
| WBGene00021888 | manf-1 | 0,84 |
| WBGene00020517 | hpo-8 | 0,84 |
| WBGene00004014 | phb-1 | 0,84 |
| WBGene00011638 | ostb-1 | 0,84 |
| WBGene00009829 | tmed-10 | 0,84 |
| WBGene00011734 | mlc-5 | 0,84 |
| WBGene00013025 | vha-13 | 0,84 |
| WBGene00000379 | cct-4 | 0,84 |
| WBGene00021420 | trap-3 | 0,84 |
| WBGene00004807 | skr-1 | 0,84 |
| WBGene00011510 | pdha-1 | 0,84 |
| WBGene00012434 | Y11D7A.10 | 0,84 |
| WBGene00004387 | rnp-4 | 0,84 |
| WBGene00018532 | pmpl-1 | 0,84 |
| WBGene00020507 | vha-15 | 0,83 |
| WBGene00020246 | clic-1 | 0,83 |
| WBGene00016790 | C49H3.3 | 0,83 |
| WBGene00010174 | F56H9.2 | 0,83 |
| WBGene00007185 | nlp-36 | 0,83 |
| WBGene00010579 | K05C4.2 | 0,83 |
| WBGene00002363 | let-92 | 0,83 |
| WBGene00004915 | snr-2 | 0,83 |
| WBGene00011059 | R06C1.4 | 0,83 |
| WBGene00019693 | ostd-1 | 0,83 |
| WBGene00001233 | eif-3.K | 0,83 |
| WBGene00001915 | his-41 | 0,83 |
| WBGene00000170 | aqp-2 | 0,83 |
| WBGene00000835 | cuc-1 | 0,83 |
| WBGene00022678 | sar-1 | 0,83 |
| WBGene00003947 | pbs-1 | 0,83 |
| WBGene00014204 | ccdc-47 | 0,83 |
| WBGene00009915 | F52A8.1 | 0,83 |

| | | |
|---|---|---|
| WBGene00009812 | suca-1 | 0,83 |
| WBGene00002074 | ima-3 | 0,83 |
| WBGene00021844 | sec-11 | 0,83 |
| WBGene00001030 | dnj-12 | 0,82 |
| WBGene00013081 | txt-14 | 0,82 |
| WBGene00001229 | eif-3.F | 0,82 |
| WBGene00006764 | unc-27 | 0,82 |
| WBGene00022114 | Y71F9AL.9 | 0,82 |
| WBGene00001503 | fum-1 | 0,82 |
| WBGene00003587 | ned-8 | 0,82 |
| WBGene00020950 | dlst-1 | 0,82 |
| WBGene00004437 | rsl-24D1 | 0,82 |
| WBGene00001393 | fat-1 | 0,82 |
| WBGene00000041 | aco-2 | 0,82 |
| WBGene00022122 | trap-1 | 0,82 |
| WBGene00006726 | ubl-5 | 0,82 |
| WBGene00006754 | unc-15 | 0,82 |
| WBGene00003924 | pas-3 | 0,82 |
| WBGene00001155 | ech-6 | 0,82 |
| WBGene00000214 | asp-1 | 0,82 |
| WBGene00001239 | elo-1 | 0,82 |
| WBGene00022793 | ZK686.3 | 0,82 |
| WBGene00003926 | pas-5 | 0,82 |
| WBGene00001088 | dpy-30 | 0,82 |
| WBGene00004266 | rab-1 | 0,81 |
| WBGene00013569 | bath-36 | 0,81 |
| WBGene00008506 | tkt-1 | 0,81 |
| WBGene00008149 | pyp-1 | 0,81 |
| WBGene00017864 | pcca-1 | 0,81 |
| WBGene00012382 | ttr-16 | 0,81 |
| WBGene00004798 | sip-1 | 0,81 |
| WBGene00002981 | lgg-2 | 0,81 |
| WBGene00018602 | F48D6.4 | 0,81 |
| WBGene00018701 | pccb-1 | 0,81 |
| WBGene00021133 | tomm-22 | 0,81 |
| WBGene00011869 | dod-6 | 0,81 |
| WBGene00021883 | Y54G2A.18 | 0,81 |
| WBGene00002054 | ifb-2 | 0,81 |
| WBGene00007824 | dlat-2 | 0,81 |
| WBGene00002891 | let-767 | 0,81 |
| WBGene00003162 | mdh-2 | 0,81 |
| WBGene00019947 | htz-1 | 0,81 |
| WBGene00007463 | vgln-1 | 0,81 |
| WBGene00012484 | car-1 | 0,81 |
| WBGene00001682 | gpc-2 | 0,81 |
| WBGene00002064 | iff-1 | 0,81 |

| | | |
|---|---|---|
| WBGene00001604 | gln-3 | 0,81 |
| WBGene00011038 | R05H5.3 | 0,81 |
| WBGene00020808 | clik-1 | 0,81 |
| WBGene00021331 | glrx-10 | 0,81 |
| WBGene00003148 | mbf-1 | 0,81 |
| WBGene00012768 | eef-1B.2 | 0,80 |
| WBGene00004990 | spp-5 | 0,80 |
| WBGene00009952 | acaa-2 | 0,80 |
| WBGene00014454 | MTCE.7 | 0,80 |
| WBGene00000181 | ard-1 | 0,80 |
| WBGene00001794 | gta-1 | 0,80 |
| WBGene00003063 | lpd-7 | 0,80 |
| WBGene00012471 | nosr-1 | 0,80 |
| WBGene00004736 | sca-1 | 0,80 |
| WBGene00004999 | spp-14 | 0,80 |
| WBGene00004467 | rpn-11 | 0,80 |
| WBGene00010993 | vha-20 | 0,80 |
| WBGene00006706 | ubc-9 | 0,80 |
| WBGene00004463 | rpn-7 | 0,79 |
| WBGene00001094 | dars-1 | 0,79 |
| WBGene00006794 | unc-60 | 0,79 |
| WBGene00016981 | rpn-13 | 0,79 |
| WBGene00003052 | lmn-1 | 0,79 |
| WBGene00019333 | moma-1 | 0,79 |
| WBGene00010266 | dct-18 | 0,79 |
| WBGene00003955 | pcn-1 | 0,79 |
| WBGene00019893 | sgt-1 | 0,79 |
| WBGene00016195 | erd-2.2 | 0,79 |
| WBGene00001909 | his-35 | 0,79 |
| WBGene00003951 | pbs-5 | 0,79 |
| WBGene00000877 | cyn-1 | 0,79 |
| WBGene00016630 | acer-1 | 0,79 |
| WBGene00007848 | cytb-5.1 | 0,79 |
| WBGene00021486 | lbp-9 | 0,79 |
| WBGene00000273 | brp-1 | 0,79 |
| WBGene00011977 | T24B8.3 | 0,78 |
| WBGene00015687 | chdp-1 | 0,78 |
| WBGene00016496 | C37C3.2 | 0,78 |
| WBGene00004320 | rbx-1 | 0,78 |
| WBGene00017347 | F10E7.5 | 0,78 |
| WBGene00003927 | pas-6 | 0,78 |
| WBGene00016653 | ssb-1 | 0,78 |
| WBGene00000381 | cct-6 | 0,78 |
| WBGene00010560 | eif-2beta | 0,78 |
| WBGene00010794 | dld-1 | 0,78 |
| WBGene00010730 | ensa-1 | 0,78 |

| | | |
|---|---|---|
| WBGene00004357 | rho-1 | 0,78 |
| WBGene00002077 | imb-3 | 0,78 |
| WBGene00009092 | tomm-20 | 0,78 |
| WBGene00009187 | etfa-1 | 0,78 |
| WBGene00018271 | ctsa-1.1 | 0,78 |
| WBGene00009918 | gcsh-2 | 0,77 |
| WBGene00010639 | K07F5.15 | 0,77 |
| WBGene00000219 | asp-6 | 0,77 |
| WBGene00010942 | chch-3 | 0,77 |
| WBGene00004305 | ran-4 | 0,77 |
| WBGene00016435 | C35B1.5 | 0,77 |
| WBGene00000216 | asp-3 | 0,77 |
| WBGene00007355 | rpb-6 | 0,77 |
| WBGene00000786 | cpr-6 | 0,77 |
| WBGene00021952 | vha-19 | 0,77 |
| WBGene00019679 | spcs-3 | 0,77 |
| WBGene00020738 | T23F2.5 | 0,77 |
| WBGene00004931 | sod-2 | 0,77 |
| WBGene00002025 | hsp-60 | 0,77 |
| WBGene00010075 | F55A11.1 | 0,77 |
| WBGene00000218 | asp-5 | 0,77 |
| WBGene00011558 | ostf-4 | 0,77 |
| WBGene00007696 | tram-1 | 0,76 |
| WBGene00015778 | got-2.2 | 0,76 |
| WBGene00017119 | timm-17B.1 | 0,76 |
| WBGene00022354 | Y82E9BR.22 | 0,76 |
| WBGene00022497 | Y119D3B.21 | 0,76 |
| WBGene00001747 | gsp-1 | 0,76 |
| WBGene00001898 | his-24 | 0,76 |
| WBGene00001333 | erm-1 | 0,76 |
| WBGene00012140 | dap-1 | 0,76 |
| WBGene00003588 | nex-1 | 0,76 |
| WBGene00001426 | fkb-1 | 0,76 |
| WBGene00000378 | cct-2 | 0,76 |
| WBGene00009082 | dlat-1 | 0,76 |
| WBGene00009995 | spig-9 | 0,76 |
| WBGene00006715 | ubc-20 | 0,76 |
| WBGene00014472 | MTCE.33 | 0,75 |
| WBGene00014022 | famh-136 | 0,75 |
| WBGene00004460 | rpn-3 | 0,75 |
| WBGene00004788 | sft-4 | 0,75 |
| WBGene00004505 | rpt-5 | 0,75 |
| WBGene00005663 | sars-1 | 0,75 |
| WBGene00000390 | cdc-42 | 0,75 |
| WBGene00001648 | goa-1 | 0,75 |
| WBGene00008852 | ubql-1 | 0,75 |

| | | |
|---|---|---|
| WBGene00003080 | lsm-6 | 0,75 |
| WBGene00007107 | pfd-4 | 0,75 |
| WBGene00019543 | cif-1 | 0,75 |
| WBGene00019719 | M01H9.3 | 0,75 |
| WBGene00020437 | stt-3 | 0,75 |
| WBGene00011856 | nspg-7.1 | 0,75 |
| WBGene00006920 | vha-11 | 0,74 |
| WBGene00007350 | sucl-1 | 0,74 |
| WBGene00004462 | rpn-6.1 | 0,74 |
| WBGene00005018 | sqt-3 | 0,74 |
| WBGene00003564 | ncs-2 | 0,74 |
| WBGene00017070 | praf-3 | 0,74 |
| WBGene00010677 | gtbp-1 | 0,74 |
| WBGene00009394 | clec-63 | 0,74 |
| WBGene00001679 | gpb-1 | 0,74 |
| WBGene00001244 | elo-6 | 0,74 |
| WBGene00016316 | C32D5.8 | 0,73 |
| WBGene00006912 | vha-3 | 0,73 |
| WBGene00001051 | cks-1 | 0,73 |
| WBGene00003891 | osm-11 | 0,73 |
| WBGene00019017 | bcf-1 | 0,73 |
| WBGene00012615 | dct-16 | 0,73 |
| WBGene00014109 | chpf-1 | 0,73 |
| WBGene00000276 | byn-1 | 0,73 |
| WBGene00010317 | idh-1 | 0,73 |
| WBGene00004152 | pqn-70 | 0,73 |
| WBGene00000293 | cap-2 | 0,73 |
| WBGene00000928 | dbd-2 | 0,73 |
| WBGene00004138 | pqn-53 | 0,72 |
| WBGene00001386 | far-2 | 0,72 |
| WBGene00021466 | eif-2gamma | 0,72 |
| WBGene00012739 | Y40B1B.7 | 0,72 |
| WBGene00006452 | heh-1 | 0,72 |
| WBGene00016961 | vps-32.1 | 0,72 |
| WBGene00004699 | rsp-2 | 0,72 |
| WBGene00013200 | Y54E5A.5 | 0,72 |
| WBGene00001976 | hmg-11 | 0,72 |
| WBGene00002015 | hsp-16.1 | 0,72 |
| WBGene00002017 | hsp-16.11 | 0,72 |
| WBGene00020374 | serp-1.2 | 0,71 |
| WBGene00010539 | ttr-2 | 0,71 |
| WBGene00000217 | asp-4 | 0,71 |
| WBGene00006699 | uba-1 | 0,71 |
| WBGene00016621 | arch-1 | 0,71 |
| WBGene00020391 | cct-7 | 0,71 |
| WBGene00007000 | tufm-1 | 0,71 |

| | | |
|---|---|---|
| WBGene00044784 | F13H10.8 | 0,71 |
| WBGene00015232 | B0511.6 | 0,71 |
| WBGene00003773 | nlt-1 | 0,71 |
| WBGene00008205 | sams-1 | 0,71 |
| WBGene00019003 | tmed-3 | 0,71 |
| WBGene00019759 | M03F4.6 | 0,71 |
| WBGene00020347 | ech-1.2 | 0,71 |
| WBGene00000822 | csq-1 | 0,71 |
| WBGene00000149 | apl-1 | 0,71 |
| WBGene00009583 | aagr-3 | 0,71 |
| WBGene00010244 | F58D5.5 | 0,71 |
| WBGene00004501 | rpt-1 | 0,71 |
| WBGene00007836 | phdh-1 | 0,71 |
| WBGene00002253 | lbp-1 | 0,71 |
| WBGene00011474 | aldo-1 | 0,71 |
| WBGene00013307 | melo-1 | 0,70 |
| WBGene00008764 | dylt-1 | 0,70 |
| WBGene00002061 | ife-3 | 0,70 |
| WBGene00044324 | ufm-1 | 0,70 |
| WBGene00001394 | fat-2 | 0,70 |
| WBGene00017852 | F27C1.2 | 0,70 |
| WBGene00001073 | dpy-11 | 0,70 |
| WBGene00002078 | xpo-1 | 0,70 |
| WBGene00021348 | moag-4 | 0,70 |
| WBGene00006708 | ubc-13 | 0,70 |
| WBGene00019780 | M60.4 | 0,70 |
| WBGene00010783 | mrpl-36 | 0,69 |
| WBGene00016235 | C29H12.2 | 0,69 |
| WBGene00011383 | T02E9.5 | 0,69 |
| WBGene00194986 | pigb-1 | 0,69 |
| WBGene00019760 | calu-1 | 0,69 |
| WBGene00000192 | arl-8 | 0,69 |
| WBGene00010479 | tass-1 | 0,69 |
| WBGene00021068 | W06H8.6 | 0,69 |
| WBGene00004502 | rpt-2 | 0,69 |
| WBGene00008860 | romo-1 | 0,68 |
| WBGene00002000 | hrpr-1 | 0,68 |
| WBGene00017088 | akir-1 | 0,68 |
| WBGene00019510 | nucl-1 | 0,68 |
| WBGene00016943 | acdh-1 | 0,68 |
| WBGene00006572 | tin-9.1 | 0,68 |
| WBGene00021441 | Y39A3CL.3 | 0,68 |
| WBGene00017970 | F32A5.4 | 0,68 |
| WBGene00012885 | iscu-1 | 0,68 |
| WBGene00020268 | hpo-19 | 0,68 |
| WBGene00001999 | hrpa-1 | 0,68 |

| | | |
|---|---|---|
| WBGene00022856 | cth-2 | 0,68 |
| WBGene00001396 | fat-4 | 0,68 |
| WBGene00013263 | txdc-12.1 | 0,68 |
| WBGene00010564 | ath-1 | 0,68 |
| WBGene00009440 | idhg-1 | 0,68 |
| WBGene00008741 | ctsa-1.2 | 0,68 |
| WBGene00003922 | pas-1 | 0,68 |
| WBGene00017734 | etfb-1 | 0,68 |
| WBGene00002238 | kars-1 | 0,67 |
| WBGene00007142 | ttr-18 | 0,67 |
| WBGene00011730 | drr-2 | 0,67 |
| WBGene00010204 | cpr-9 | 0,67 |
| WBGene00004464 | rpn-8 | 0,67 |
| WBGene00007352 | cdc-48.1 | 0,67 |
| WBGene00002269 | lec-6 | 0,67 |
| WBGene00004504 | rpt-4 | 0,67 |
| WBGene00002020 | hsp-16.49 | 0,67 |
| WBGene00002019 | hsp-16.48 | 0,67 |
| WBGene00016749 | C48E7.1 | 0,67 |
| WBGene00013360 | tmed-4 | 0,67 |
| WBGene00009045 | F22B5.10 | 0,67 |
| WBGene00009783 | rer-1 | 0,67 |
| WBGene00011648 | cni-1 | 0,67 |
| WBGene00013577 | mmad-1 | 0,67 |
| WBGene00020366 | acdh-10 | 0,67 |
| WBGene00003925 | pas-4 | 0,67 |
| WBGene00020038 | R12E2.13 | 0,66 |
| WBGene00009367 | F33H2.3 | 0,66 |
| WBGene00000479 | cgh-1 | 0,66 |
| WBGene00019607 | K10B2.4 | 0,66 |
| WBGene00011735 | hip-1 | 0,66 |
| WBGene00001230 | eif-3.G | 0,66 |
| WBGene00011481 | imp-2 | 0,66 |
| WBGene00022492 | dss-1 | 0,66 |
| WBGene00010049 | F54D5.3 | 0,66 |
| WBGene00000122 | aly-3 | 0,66 |
| WBGene00008446 | E01G4.3 | 0,66 |
| WBGene00022734 | ZK418.5 | 0,66 |
| WBGene00015803 | C15H9.9 | 0,66 |
| WBGene00001858 | hil-7 | 0,66 |
| WBGene00001076 | dpy-17 | 0,66 |
| WBGene00011304 | mnk-1 | 0,66 |
| WBGene00001404 | fbp-1 | 0,65 |
| WBGene00003163 | mdl-1 | 0,65 |
| WBGene00003795 | npp-9 | 0,65 |
| WBGene00006702 | ubc-3 | 0,65 |

| | | |
|---|---|---|
| WBGene00020679 | ogdh-1 | 0,65 |
| WBGene00003786 | npa-1 | 0,65 |
| WBGene00002273 | lec-10 | 0,65 |
| WBGene00012015 | T25B9.9 | 0,65 |
| WBGene00022647 | slc-17.1 | 0,64 |
| WBGene00006720 | ubc-25 | 0,64 |
| WBGene00002059 | ife-1 | 0,64 |
| WBGene00018764 | azin-1 | 0,64 |
| WBGene00007449 | C08F8.9 | 0,64 |
| WBGene00001075 | dpy-14 | 0,64 |
| WBGene00002008 | hsp-4 | 0,64 |
| WBGene00020662 | T21H3.1 | 0,64 |
| WBGene00020107 | prps-1 | 0,64 |
| WBGene00018008 | vpr-1 | 0,64 |
| WBGene00012907 | cpt-1 | 0,64 |
| WBGene00006915 | vha-6 | 0,64 |
| WBGene00017490 | pud-2.1 | 0,63 |
| WBGene00017500 | pud-2.2 | 0,63 |
| WBGene00017993 | cec-5 | 0,63 |
| WBGene00013378 | emc-3 | 0,63 |
| WBGene00001993 | hpd-1 | 0,62 |
| WBGene00000107 | alh-1 | 0,62 |
| WBGene00017311 | F09G2.2 | 0,62 |
| WBGene00001234 | eif-6 | 0,62 |
| WBGene00008572 | F08B12.4 | 0,62 |
| WBGene00020812 | acdh-7 | 0,62 |
| WBGene00021548 | trx-4 | 0,62 |
| WBGene00021325 | nspg-13 | 0,61 |
| WBGene00020115 | mboa-6 | 0,61 |
| WBGene00015125 | hadb-1 | 0,61 |
| WBGene00004466 | rpn-10 | 0,61 |
| WBGene00009219 | dpm-3 | 0,61 |
| WBGene00017591 | nspg-10 | 0,61 |
| WBGene00019354 | K03B4.2 | 0,61 |
| WBGene00015413 | pdhb-1 | 0,60 |
| WBGene00016011 | C23G10.2 | 0,60 |
| WBGene00006573 | tin-10 | 0,60 |
| WBGene00010605 | kola-4 | 0,60 |
| WBGene00010778 | gpdh-2 | 0,59 |
| WBGene00021956 | Y57E12AL.1 | 0,59 |
| WBGene00001226 | eif-3.C | 0,58 |
| WBGene00001209 | egl-45 | 0,58 |
| WBGene00008398 | D2005.3 | 0,57 |
| WBGene00008053 | cdc-48.2 | 0,53 |
