## Supplementary material for "Confidence-aware learning for transcriptome-based prediction of OXPHOS genes in *Caenorhabditis elegans* under incomplete functional annotation": Table S6

| Gene | Public.Name | Stage | Type.of.gene | Probability |
|---|---|---|---|---|
| WBGene00000207 | asb-2 | Embryo, Adult | High-confidence positive | |
| WBGene00000210 | asg-2 | Embryo, Adult | High-confidence positive | |
| WBGene00000229 | atp-2 | Embryo, Adult | High-confidence positive | |
| WBGene00000230 | atp-3 | Embryo, Adult | High-confidence positive | |
| WBGene00000869 | cyc-1 | Embryo, Adult | High-confidence positive | |
| WBGene00003061 | lpd-5 | Embryo, Adult | High-confidence positive | |
| WBGene00003831 | nuo-1 | Embryo, Adult | High-confidence positive | |
| WBGene00006519 | cox-6A | Embryo, Adult | High-confidence positive | |
| WBGene00007385 | atp-5 | Embryo, Adult | High-confidence positive | |
| WBGene00008414 | ndub-7 | Embryo, Adult | High-confidence positive | |
| WBGene00009992 | nduv-2 | Embryo, Adult | High-confidence positive | |
| WBGene00010419 | atp-1 | Embryo, Adult | High-confidence positive | |
| WBGene00011015 | R04F11.2 | Embryo, Adult | High-confidence positive | |
| WBGene00011273 | R53.4 | Embryo, Adult | High-confidence positive | |
| WBGene00012094 | T27E9.2 | Embryo, Adult | High-confidence positive | |
| WBGene00012166 | nuo-6 | Embryo, Adult | High-confidence positive | |
| WBGene00012376 | nduf-7 | Embryo, Adult | High-confidence positive | |
| WBGene00015810 | C16A3.5 | Embryo, Adult | High-confidence positive | |
| WBGene00016118 | ndub-5 | Embryo, Adult | High-confidence positive | |
| WBGene00017121 | cyc-2.1 | Embryo, Adult | High-confidence positive | |
| WBGene00017925 | F29B9.11 | Embryo, Adult | High-confidence positive | |
| WBGene00018963 | ucr-1 | Embryo, Adult | High-confidence positive | |
| WBGene00019007 | ucr-11 | Embryo, Adult | High-confidence positive | |
| WBGene00019061 | F58F12.1 | Embryo, Adult | High-confidence positive | |
| WBGene00020275 | atp-4 | Embryo, Adult | High-confidence positive | |
| WBGene00020417 | nuo-2 | Embryo, Adult | High-confidence positive | |
| WBGene00020636 | ndus-8 | Embryo, Adult | High-confidence positive | |
| WBGene00021800 | nduf-9 | Embryo, Adult | High-confidence positive | |
| WBGene00022089 | Y69A2AR.18 | Embryo, Adult | High-confidence positive | |
| WBGene00022336 | Y82E9BR.3 | Embryo, Adult | High-confidence positive | |
| WBGene00000829 | ctb-1 | Embryo | High-confidence positive | |
| WBGene00010326 | ndub-10 | Embryo | High-confidence positive | |
| WBGene00010957 | nduo-6 | Embryo | High-confidence positive | |
| WBGene00010959 | nduo-1 | Embryo | High-confidence positive | |
| WBGene00010960 | atp-6 | Embryo | High-confidence positive | |
| WBGene00010961 | nduo-2 | Embryo | High-confidence positive | |
| WBGene00010962 | ctc-3 | Embryo | High-confidence positive | |
| WBGene00010963 | nduo-4 | Embryo | High-confidence positive | |
| WBGene00010964 | ctc-1 | Embryo | High-confidence positive | |
| WBGene00010965 | ctc-2 | Embryo | High-confidence positive | |
| WBGene00010967 | nduo-5 | Embryo | High-confidence positive | |
| WBGene00013804 | Y116A8C.27 | Embryo | High-confidence positive | |
| WBGene00018151 | ndab-2 | Embryo | High-confidence positive | |
| WBGene00021562 | nuo-5 | Embryo | High-confidence positive | |
| WBGene00000371 | cox-5B | Adult | High-confidence positive | |
| WBGene00002162 | isp-1 | Adult | High-confidence positive | |
| WBGene00003225 | mev-1 | Adult | High-confidence positive | |

| | | | | |
|---|---|---|---|---|
| WBGene00006433 | sdhb-1 | Adult | High-confidence positive | |
| WBGene00007192 | nduf-11 | Adult | High-confidence positive | |
| WBGene00007684 | ndub-3 | Adult | High-confidence positive | |
| WBGene00007880 | ndua-5 | Adult | High-confidence positive | |
| WBGene00009051 | nduf-6 | Adult | High-confidence positive | |
| WBGene00009161 | cox-7C | Adult | High-confidence positive | |
| WBGene00009294 | ndua-1 | Adult | High-confidence positive | |
| WBGene00009454 | F36A2.7 | Adult | High-confidence positive | |
| WBGene00009712 | ndub-2 | Adult | High-confidence positive | |
| WBGene00012354 | cox-4 | Adult | High-confidence positive | |
| WBGene00012553 | cox-5A | Adult | High-confidence positive | |
| WBGene00013094 | ndub-8 | Adult | High-confidence positive | |
| WBGene00013308 | nuo-3 | Adult | High-confidence positive | |
| WBGene00013406 | ndua-2 | Adult | High-confidence positive | |
| WBGene00014086 | ndub-6 | Adult | High-confidence positive | |
| WBGene00015755 | C14B9.10 | Adult | High-confidence positive | |
| WBGene00016393 | ndua-13 | Adult | High-confidence positive | |
| WBGene00017926 | cox-6C | Adult | High-confidence positive | |
| WBGene00018361 | ndub-11 | Adult | High-confidence positive | |
| WBGene00020181 | T02H6.11 | Adult | High-confidence positive | |
| WBGene00021849 | ndua-8 | Adult | High-confidence positive | |
| WBGene00022169 | nduc-2 | Adult | High-confidence positive | |
| WBGene00022170 | cox-6B | Adult | High-confidence positive | |
| WBGene00014095 | gdh-1 | Adult | Predicted | 0,901 |
| WBGene00004470 | rps-3A | Embryo | Predicted | 0,901 |
| WBGene00004418 | rpl-7 | Embryo | Predicted | 0,902 |
| WBGene00004449 | rpl-35 | Embryo | Predicted | 0,902 |
| WBGene00004423 | rpl-11.2 | Embryo | Predicted | 0,903 |
| WBGene00004494 | rps-25 | Embryo | Predicted | 0,903 |
| WBGene00001423 | fib-1 | Embryo | Predicted | 0,905 |
| WBGene00009772 | ztf-7 | Embryo | Predicted | 0,905 |
| WBGene00004469 | rpsa-1 | Embryo | Predicted | 0,906 |
| WBGene00001167 | eef-2 | Embryo | Predicted | 0,907 |
| WBGene00004431 | rpl-19 | Embryo | Predicted | 0,907 |
| WBGene00004409 | rplp-1 | Embryo | Predicted | 0,907 |
| WBGene00004421 | rpl-10L | Embryo | Predicted | 0,907 |
| WBGene00004483 | rps-14 | Embryo | Predicted | 0,907 |
| WBGene00004480 | rps-11 | Embryo | Predicted | 0,908 |
| WBGene00013766 | prmt-1 | Embryo | Predicted | 0,908 |
| WBGene00004415 | rpl-4 | Embryo | Predicted | 0,908 |
| WBGene00004429 | rpl-17 | Embryo | Predicted | 0,909 |
| WBGene00004441 | rpl-27 | Embryo | Predicted | 0,909 |
| WBGene00004444 | rpl-30 | Embryo | Predicted | 0,909 |
| WBGene00010478 | dkc-1 | Embryo | Predicted | 0,910 |
| WBGene00004436 | rpl-24 | Embryo | Predicted | 0,911 |
| WBGene00004487 | rps-18 | Embryo | Predicted | 0,911 |
| WBGene00021350 | rpl-27A | Embryo | Predicted | 0,911 |

| | | | | |
|---|---|---|---|---|
| WBGene00016493 | rplp-2.1 | Embryo | Predicted | 0,911 |
| WBGene00004426 | rpl-14 | Embryo | Predicted | 0,912 |
| WBGene00004419 | rpl-7A | Embryo | Predicted | 0,912 |
| WBGene00004408 | rplp-0 | Embryo | Predicted | 0,912 |
| WBGene00004472 | rps-3 | Embryo | Predicted | 0,912 |
| WBGene00006725 | rps-27A | Embryo | Predicted | 0,913 |
| WBGene00012344 | ola-1 | Embryo | Predicted | 0,913 |
| WBGene00004447 | rpl-35A | Embryo | Predicted | 0,913 |
| WBGene00006728 | ubq-2 | Embryo | Predicted | 0,913 |
| WBGene00007630 | har-1 | Embryo, Adult | Predicted | 0,914 |
| WBGene00001169 | eef-1A.2 | Embryo | Predicted | 0,914 |
| WBGene00004428 | rpl-13A | Embryo | Predicted | 0,915 |
| WBGene00017984 | gmpr-1 | Embryo, Adult | Predicted | 0,915 |
| WBGene00004478 | rps-9 | Embryo | Predicted | 0,916 |
| WBGene00019680 | K12H4.5 | Adult | Predicted | 0,917 |
| WBGene00004410 | rplp-2.2 | Embryo | Predicted | 0,919 |
| WBGene00004451 | rpl-37.2 | Embryo | Predicted | 0,919 |
| WBGene00010896 | snu-13 | Embryo | Predicted | 0,919 |
| WBGene00021248 | Y22D7AL.10 | Embryo | Predicted | 0,919 |
| WBGene00017075 | nap-1 | Embryo | Predicted | 0,920 |
| WBGene00003065 | lpd-9 | Adult | Predicted | 0,921 |
| WBGene00004493 | rps-24 | Embryo | Predicted | 0,921 |
| WBGene00004492 | rps-23 | Embryo | Predicted | 0,921 |
| WBGene00004488 | rps-19 | Embryo | Predicted | 0,921 |
| WBGene00004479 | rps-10 | Embryo | Predicted | 0,922 |
| WBGene00004456 | rpl-37A | Embryo | Predicted | 0,922 |
| WBGene00004302 | ran-1 | Embryo | Predicted | 0,922 |
| WBGene00004434 | rpl-22 | Embryo | Predicted | 0,923 |
| WBGene00004489 | rps-20 | Embryo | Predicted | 0,923 |
| WBGene00004484 | rps-15 | Embryo | Predicted | 0,923 |
| WBGene00012179 | rpl-37.1 | Embryo | Predicted | 0,924 |
| WBGene00004439 | rpl-23A.2 | Embryo | Predicted | 0,924 |
| WBGene00004450 | rpl-36 | Embryo | Predicted | 0,925 |
| WBGene00004442 | rpl-28 | Embryo | Predicted | 0,925 |
| WBGene00007708 | nola-3 | Embryo | Predicted | 0,926 |
| WBGene00004474 | rps-5 | Embryo | Predicted | 0,927 |
| WBGene00077526 | C25A1.16 | Embryo | Predicted | 0,927 |
| WBGene00012602 | Y38E10A.24 | Adult | Predicted | 0,930 |
| WBGene00001168 | eef-1A.1 | Embryo | Predicted | 0,931 |
| WBGene00004471 | rps-2 | Embryo | Predicted | 0,933 |
| WBGene00004424 | rpl-12 | Embryo | Predicted | 0,934 |
| WBGene00009122 | tct-1 | Embryo | Predicted | 0,935 |
| WBGene00003902 | pab-1 | Embryo | Predicted | 0,936 |
| WBGene00022235 | sqd-1 | Embryo | Predicted | 0,937 |
| WBGene00015248 | mai-2 | Embryo, Adult | Predicted | 0,938 |
| WBGene00006439 | ant-1.1 | Embryo, Adult | Predicted | 0,940 |
| WBGene00006924 | vig-1 | Embryo | Predicted | 0,944 |

| | | | | |
|---|---|---|---|---|
| WBGene00004435 | rpl-23 | Embryo | Predicted | 0,945 |
| WBGene00019537 | K08D12.3 | Embryo | Predicted | 0,946 |
| WBGene00018846 | eef-1B.1 | Embryo | Predicted | 0,946 |
| WBGene00008920 | eef-1G | Embryo | Predicted | 0,948 |
| WBGene00011156 | rbm-3.2 | Embryo | Predicted | 0,949 |
| WBGene00018519 | F46H5.3 | Adult | Predicted | 0,952 |
| WBGene00009119 | ndk-1 | Embryo | Predicted | 0,953 |
| WBGene00008505 | F01G4.6 | Embryo | Predicted | 0,957 |
| WBGene00002045 | icd-1 | Embryo | Predicted | 0,957 |
| WBGene00000833 | cts-1 | Embryo | Predicted | 0,958 |
| WBGene00013463 | kdp-1 | Adult | Predicted | 0,960 |
| WBGene00000263 | F23H11.5 | Adult | Predicted | 0,961 |
| WBGene00008262 | ril-1 | Embryo, Adult | Predicted | 0,962 |
| WBGene00009688 | F44E5.1 | Adult | Predicted | 0,965 |
| WBGene00001209 | egl-45 | Embryo | | 0,575 |
| WBGene00001226 | eif-3.C | Embryo | | 0,581 |
| WBGene00006573 | tin-10 | Embryo | | 0,603 |
| WBGene00015413 | pdhb-1 | Adult | | 0,604 |
| WBGene00001234 | eif-6 | Embryo | | 0,620 |
| WBGene00011730 | drr-2 | Embryo | | 0,672 |
| WBGene00002238 | kars-1 | Embryo | | 0,675 |
| WBGene00017734 | etfb-1 | Embryo | | 0,675 |
| WBGene00020268 | hpo-19 | Embryo | | 0,680 |
| WBGene00019510 | nucl-1 | Embryo | | 0,682 |
| WBGene00008860 | romo-1 | Embryo | | 0,684 |
| WBGene00001394 | fat-2 | Embryo | | 0,701 |
| WBGene00015232 | B0511.6 | Embryo | | 0,710 |
| WBGene00007000 | tufm-1 | Embryo | | 0,710 |
| WBGene00000276 | byn-1 | Embryo | | 0,730 |
| WBGene00019543 | cif-1 | Embryo | | 0,748 |
| WBGene00005663 | sars-1 | Embryo | | 0,752 |
| WBGene00009082 | dlat-1 | Adult | | 0,757 |
| WBGene00002025 | hsp-60 | Embryo | | 0,768 |
| WBGene00007355 | rpb-6 | Embryo | | 0,771 |
| WBGene00010942 | chch-3 | Adult | | 0,773 |
| WBGene00010639 | K07F5.15 | Embryo | | 0,775 |
| WBGene00009092 | tomm-20 | Embryo | | 0,776 |
| WBGene00002077 | imb-3 | Embryo | | 0,776 |
| WBGene00016653 | ssb-1 | Embryo | | 0,780 |
| WBGene00017347 | F10E7.5 | Embryo | | 0,781 |
| WBGene00000877 | cyn-1 | Embryo | | 0,789 |
| WBGene00003063 | lpd-7 | Embryo | | 0,802 |
| WBGene00012768 | eef-1B.2 | Embryo | | 0,805 |
| WBGene00003162 | mdh-2 | Adult | | 0,809 |
| WBGene00007824 | dlat-2 | Adult | | 0,809 |
| WBGene00021133 | tomm-22 | Embryo | | 0,810 |
| WBGene00000041 | aco-2 | Embryo | | 0,822 |

| | | | |
|---|---|---|---|
| WBGene00004437 | rsl-24D1 | Embryo | 0,822 |
| WBGene00011059 | R06C1.4 | Embryo | 0,830 |
| WBGene00016790 | C49H3.3 | Embryo | 0,834 |
| WBGene00004014 | phb-1 | Embryo | 0,839 |
| WBGene00004015 | phb-2 | Embryo | 0,842 |
| WBGene00021024 | W04C9.2 | Adult | 0,842 |
| WBGene00010303 | cri-3 | Embryo | 0,843 |
| WBGene00012964 | Y48A6B.3 | Embryo | 0,849 |
| WBGene00077771 | tspo-1 | Adult | 0,849 |
| WBGene00020915 | nol-58 | Embryo | 0,849 |
| WBGene00002010 | hsp-6 | Embryo | 0,851 |
| WBGene00009664 | idha-1 | Adult | 0,852 |
| WBGene00020297 | nog-1 | Embryo | 0,855 |
| WBGene00021088 | W08E12.7 | Embryo | 0,860 |
| WBGene00022053 | cisd-3.2 | Embryo, Adult | 0,861 |
| WBGene00020931 | cytb-5.2 | Embryo | 0,861 |
| WBGene00004413 | rpl-8 | Embryo | 0,866 |
| WBGene00000941 | ddp-1 | Embryo | 0,872 |
| WBGene00004497 | rps-28 | Embryo | 0,873 |
| WBGene00004443 | rpl-29 | Embryo | 0,874 |
| WBGene00002065 | iff-2 | Embryo | 0,875 |
| WBGene00004495 | rps-26 | Embryo | 0,877 |
| WBGene00004453 | rpl-39 | Embryo | 0,878 |
| WBGene00004485 | rps-16 | Embryo | 0,879 |
| WBGene00012351 | W09C5.1 | Embryo | 0,881 |
| WBGene00002083 | inf-1 | Embryo | 0,881 |
| WBGene00020683 | ribo-1 | Embryo | 0,882 |
| WBGene00018784 | F54A3.5 | Adult | 0,883 |
| WBGene00012097 | abcf-2 | Embryo | 0,883 |
| WBGene00004427 | rpl-15 | Embryo | 0,884 |
| WBGene00010627 | nol-56 | Embryo | 0,885 |
| WBGene00022042 | icd-2 | Embryo | 0,886 |
| WBGene00004490 | rps-21 | Embryo | 0,887 |
| WBGene00004475 | rps-6 | Embryo | 0,887 |
| WBGene00004473 | rps-4 | Embryo | 0,887 |
| WBGene00004448 | rpl-34 | Embryo | 0,887 |
| WBGene00004454 | rpl-36A | Embryo | 0,888 |
| WBGene00004425 | rpl-13 | Embryo | 0,889 |
| WBGene00004438 | rpl-23A.1 | Embryo | 0,889 |
| WBGene00004491 | rps-15A | Embryo | 0,889 |
| WBGene00004452 | rpl-38 | Embryo | 0,890 |
| WBGene00004486 | rps-17 | Embryo | 0,892 |
| WBGene00004414 | rpl-3 | Embryo | 0,892 |
| WBGene00004481 | rps-12 | Embryo | 0,892 |
| WBGene00004416 | rpl-5 | Embryo | 0,892 |
| WBGene00004498 | rps-29 | Embryo | 0,893 |
| WBGene00004420 | rpl-9 | Embryo | 0,893 |

| | | | | |
|---|---|---|---|---|
| WBGene00010556 | rack-1 | Embryo | | 0,894 |
| WBGene00004499 | fau-1 | Embryo | | 0,895 |
| WBGene00004430 | rpl-18 | Embryo | | 0,895 |
| WBGene00004417 | rpl-6 | Embryo | | 0,895 |
| WBGene00004482 | rps-13 | Embryo | | 0,897 |
| WBGene00004440 | rpl-26 | Embryo | | 0,897 |
| WBGene00004433 | rpl-21 | Embryo | | 0,898 |
| WBGene00022783 | tomm-7 | Embryo | | 0,898 |
| WBGene00004446 | rpl-32 | Embryo | | 0,899 |
| WBGene00004412 | rpl-10a | Embryo | | 0,899 |
| WBGene00004477 | rps-8 | Embryo | | 0,900 |
| WBGene00000896 | dad-1 | Embryo | | 0,900 |
| WBGene00004445 | rpl-31 | Embryo | | 0,900 |
| WBGene00004476 | rps-7 | Embryo | | 0,900 |
| WBGene00004496 | rps-27 | Embryo | | 0,900 |
| WBGene00004432 | rpl-18A | Embryo | | 0,900 |
| WBGene00002183 | kat-1 | Embryo, Adult | NA | |
| WBGene00006937 | wah-1 | Embryo, Adult | NA | |
| WBGene00010616 | K07A1.10 | Embryo, Adult | NA | |
| WBGene00000108 | alh-2 | Embryo | NA | |
| WBGene00000209 | asg-1 | Embryo | NA | |
| WBGene00000550 | clu-1 | Embryo | NA | |
| WBGene00000761 | coq-1 | Embryo | NA | |
| WBGene00000767 | coq-8 | Embryo | NA | |
| WBGene00000931 | dao-5 | Embryo | NA | |
| WBGene00000933 | dap-3 | Embryo | NA | |
| WBGene00001039 | dnj-21 | Embryo | NA | |
| WBGene00001186 | egl-18 | Embryo | NA | |
| WBGene00001225 | eif-3.B | Embryo | NA | |
| WBGene00001227 | eif-3.D | Embryo | NA | |
| WBGene00001231 | eif-3.H | Embryo | NA | |
| WBGene00001337 | ears-1 | Embryo | NA | |
| WBGene00001435 | fkh-3 | Embryo | NA | |
| WBGene00001497 | fars-1 | Embryo | NA | |
| WBGene00001520 | gas-1 | Embryo | NA | |
| WBGene00001596 | gld-2 | Embryo | NA | |
| WBGene00001635 | gly-10 | Embryo | NA | |
| WBGene00001913 | his-39 | Embryo | NA | |
| WBGene00001945 | his-71 | Embryo | NA | |
| WBGene00002060 | ife-2 | Embryo | NA | |
| WBGene00002152 | iars-1 | Embryo | NA | |
| WBGene00002850 | let-716 | Embryo | NA | |
| WBGene00002855 | let-721 | Embryo | NA | |
| WBGene00003062 | lpd-6 | Embryo | NA | |
| WBGene00003066 | lpl-1 | Embryo | NA | |
| WBGene00003119 | mac-1 | Embryo | NA | |
| WBGene00003130 | map-2 | Embryo | NA | |

| | | | |
|---|---|---|---|
| WBGene00003415 | mars-1 | Embryo | NA |
| WBGene00003596 | ngp-1 | Embryo | NA |
| WBGene00003807 | npr-1 | Embryo | NA |
| WBGene00003815 | nars-1 | Embryo | NA |
| WBGene00003821 | nst-1 | Embryo | NA |
| WBGene00004245 | puf-9 | Embryo | NA |
| WBGene00004248 | pus-1 | Embryo | NA |
| WBGene00004727 | sax-1 | Embryo | NA |
| WBGene00004771 | sem-2 | Embryo | NA |
| WBGene00004815 | skr-9 | Embryo | NA |
| WBGene00004978 | spg-7 | Embryo | NA |
| WBGene00004981 | spl-1 | Embryo | NA |
| WBGene00005613 | srj-25 | Embryo | NA |
| WBGene00006497 | tag-151 | Embryo | NA |
| WBGene00006499 | ntl-2.2 | Embryo | NA |
| WBGene00006566 | tgt-1 | Embryo | NA |
| WBGene00006567 | tgt-2 | Embryo | NA |
| WBGene00006613 | trm-1 | Embryo | NA |
| WBGene00006617 | tars-1 | Embryo | NA |
| WBGene00006936 | glp-4 | Embryo | NA |
| WBGene00007001 | tufm-2 | Embryo | NA |
| WBGene00007194 | dph-5 | Embryo | NA |
| WBGene00007215 | oxa-1 | Embryo | NA |
| WBGene00007555 | dohh-1 | Embryo | NA |
| WBGene00007564 | mrps-22 | Embryo | NA |
| WBGene00007587 | mma-1 | Embryo | NA |
| WBGene00007616 | rpoa-12 | Embryo | NA |
| WBGene00007617 | rrsr-1 | Embryo | NA |
| WBGene00007623 | utp-11 | Embryo | NA |
| WBGene00007628 | C16C10.8 | Embryo | NA |
| WBGene00007681 | C18D11.3 | Embryo | NA |
| WBGene00007686 | tomm-40 | Embryo | NA |
| WBGene00007712 | mrpl-34 | Embryo | NA |
| WBGene00007784 | ruvb-1 | Embryo | NA |
| WBGene00007919 | cup-16 | Embryo | NA |
| WBGene00008070 | C43D7.8 | Embryo | NA |
| WBGene00008117 | gsr-1 | Embryo | NA |
| WBGene00008134 | C47B2.9 | Embryo | NA |
| WBGene00008151 | rrp-1 | Embryo | NA |
| WBGene00008155 | C47E12.12 | Embryo | NA |
| WBGene00008261 | C52G5.3 | Embryo | NA |
| WBGene00008364 | slc-25A26 | Embryo | NA |
| WBGene00008375 | D1054.8 | Embryo | NA |
| WBGene00008413 | D2030.3 | Embryo | NA |
| WBGene00008452 | mrps-5 | Embryo | NA |
| WBGene00008456 | eral-1 | Embryo | NA |
| WBGene00008459 | mrpl-53 | Embryo | NA |

| | | | |
|---|---|---|---|
| WBGene00008514 | mrpl-44 | Embryo | NA |
| WBGene00008679 | F11A5.13 | Embryo | NA |
| WBGene00008769 | mrpl-13 | Embryo | NA |
| WBGene00008781 | rpoa-2 | Embryo | NA |
| WBGene00008857 | timm-23 | Embryo | NA |
| WBGene00008948 | F19B6.1 | Embryo | NA |
| WBGene00009013 | mrps-33 | Embryo | NA |
| WBGene00009117 | F25H2.3 | Embryo | NA |
| WBGene00009128 | mrpl-54 | Embryo | NA |
| WBGene00009189 | F27D4.4 | Embryo | NA |
| WBGene00009207 | F28C6.8 | Embryo | NA |
| WBGene00009217 | spe-45 | Embryo | NA |
| WBGene00009246 | gfm-1 | Embryo | NA |
| WBGene00009248 | F29C12.6 | Embryo | NA |
| WBGene00009289 | exos-7 | Embryo | NA |
| WBGene00009330 | F32D8.5 | Embryo | NA |
| WBGene00009352 | F33A8.4 | Embryo | NA |
| WBGene00009660 | F43G6.8 | Embryo | NA |
| WBGene00009993 | F53F4.11 | Embryo | NA |
| WBGene00010015 | atad-3 | Embryo | NA |
| WBGene00010035 | F54C8.1 | Embryo | NA |
| WBGene00010094 | tsfm-1 | Embryo | NA |
| WBGene00010435 | pwp-1 | Embryo | NA |
| WBGene00010450 | metl-18 | Embryo | NA |
| WBGene00010557 | mspn-1 | Embryo | NA |
| WBGene00010582 | K05C4.5 | Embryo | NA |
| WBGene00010617 | K07A1.13 | Embryo | NA |
| WBGene00010624 | mrps-15 | Embryo | NA |
| WBGene00010638 | K07F5.14 | Embryo | NA |
| WBGene00010721 | nuaf-5 | Embryo | NA |
| WBGene00010812 | mrpl-35 | Embryo | NA |
| WBGene00010872 | plag-15 | Embryo | NA |
| WBGene00010905 | mrps-34 | Embryo | NA |
| WBGene00010909 | cisd-3.1 | Embryo | NA |
| WBGene00011043 | rbm-28 | Embryo | NA |
| WBGene00011116 | pdcd-2 | Embryo | NA |
| WBGene00011239 | pges-2 | Embryo | NA |
| WBGene00011269 | sbsp-1 | Embryo | NA |
| WBGene00011290 | R102.3 | Embryo | NA |
| WBGene00011309 | R186.8 | Embryo | NA |
| WBGene00011391 | mrps-12 | Embryo | NA |
| WBGene00011408 | nifk-1 | Embryo | NA |
| WBGene00011412 | mrpl-16 | Embryo | NA |
| WBGene00011526 | cox-15 | Embryo | NA |
| WBGene00011634 | T09A5.5 | Embryo | NA |
| WBGene00011639 | ztf-17 | Embryo | NA |
| WBGene00011679 | ucr-2.2 | Embryo | NA |

| | | | |
|---|---|---|---|
| WBGene00011857 | coa-4 | Embryo | NA |
| WBGene00011875 | cox-14 | Embryo | NA |
| WBGene00011897 | scpl-4 | Embryo | NA |
| WBGene00012030 | T25G3.3 | Embryo | NA |
| WBGene00012126 | T28D6.6 | Embryo | NA |
| WBGene00012192 | trm-61A | Embryo | NA |
| WBGene00012361 | mrpl-12 | Embryo | NA |
| WBGene00012455 | Y17D7C.1 | Embryo | NA |
| WBGene00012460 | dhps-1 | Embryo | NA |
| WBGene00012556 | mrps-10 | Embryo | NA |
| WBGene00012590 | nspe-3 | Embryo | NA |
| WBGene00012592 | Y38E10A.14 | Embryo | NA |
| WBGene00012603 | nspe-6 | Embryo | NA |
| WBGene00012645 | mrpl-22 | Embryo | NA |
| WBGene00012652 | emg-1 | Embryo | NA |
| WBGene00012676 | pro-3 | Embryo | NA |
| WBGene00012692 | nol-53 | Embryo | NA |
| WBGene00012697 | mrps-35 | Embryo | NA |
| WBGene00012714 | abce-1 | Embryo | NA |
| WBGene00012738 | eif-3.J | Embryo | NA |
| WBGene00012794 | Y43E12A.2 | Embryo | NA |
| WBGene00012830 | mrps-28 | Embryo | NA |
| WBGene00012858 | Y44F5A.1 | Embryo | NA |
| WBGene00012887 | Y45F10D.7 | Embryo | NA |
| WBGene00012978 | bop-1 | Embryo | NA |
| WBGene00012992 | mrpl-20 | Embryo | NA |
| WBGene00012999 | rpoa-1 | Embryo | NA |
| WBGene00013004 | mrpl-37 | Embryo | NA |
| WBGene00013017 | Y48E1C.4 | Embryo | NA |
| WBGene00013029 | glrx-5 | Embryo | NA |
| WBGene00013143 | Y53C12B.1 | Embryo | NA |
| WBGene00013144 | pno-1 | Embryo | NA |
| WBGene00013210 | Y54G9A.7 | Embryo | NA |
| WBGene00013236 | znf-593 | Embryo | NA |
| WBGene00013267 | Y57A10A.27 | Embryo | NA |
| WBGene00013382 | gcl-1 | Embryo | NA |
| WBGene00013435 | psme-3 | Embryo | NA |
| WBGene00013524 | Y73F8A.15 | Embryo | NA |
| WBGene00013605 | Y95D11A.1 | Embryo | NA |
| WBGene00013983 | ZK512.2 | Embryo | NA |
| WBGene00013997 | eelo-2 | Embryo | NA |
| WBGene00014120 | trmt-6 | Embryo | NA |
| WBGene00014155 | ZK930.5 | Embryo | NA |
| WBGene00014172 | clpp-1 | Embryo | NA |
| WBGene00014205 | metl-6 | Embryo | NA |
| WBGene00014224 | mrps-23 | Embryo | NA |
| WBGene00015006 | B0041.1 | Embryo | NA |

| | | | |
|---|---|---|---|
| WBGene00015021 | nfs-1 | Embryo | NA |
| WBGene00015025 | mrpl-9 | Embryo | NA |
| WBGene00015104 | B0280.9 | Embryo | NA |
| WBGene00015133 | mrpl-11 | Embryo | NA |
| WBGene00015185 | mrpl-41 | Embryo | NA |
| WBGene00015247 | B0545.4 | Embryo | NA |
| WBGene00015265 | B0563.8 | Embryo | NA |
| WBGene00015344 | nra-4 | Embryo | NA |
| WBGene00015425 | lrpr-1 | Embryo | NA |
| WBGene00015450 | C04F5.8 | Embryo | NA |
| WBGene00015461 | krr-1 | Embryo | NA |
| WBGene00015476 | C05D9.9 | Embryo | NA |
| WBGene00015487 | mrps-17 | Embryo | NA |
| WBGene00015538 | sams-3 | Embryo | NA |
| WBGene00015540 | sams-4 | Embryo | NA |
| WBGene00015811 | C16A3.6 | Embryo | NA |
| WBGene00015915 | C17G10.1 | Embryo | NA |
| WBGene00015920 | eif-3.L | Embryo | NA |
| WBGene00015941 | C18A3.3 | Embryo | NA |
| WBGene00016073 | C24H12.4 | Embryo | NA |
| WBGene00016135 | C26B9.6 | Embryo | NA |
| WBGene00016142 | mrps-18C | Embryo | NA |
| WBGene00016148 | C26F1.3 | Embryo | NA |
| WBGene00016166 | C27F2.4 | Embryo | NA |
| WBGene00016249 | mrpl-32 | Embryo | NA |
| WBGene00016412 | mrps-26 | Embryo | NA |
| WBGene00016492 | etfm-1 | Embryo | NA |
| WBGene00016589 | ltah-1.1 | Embryo | NA |
| WBGene00016607 | C43E11.9 | Embryo | NA |
| WBGene00016739 | pitr-1 | Embryo | NA |
| WBGene00016740 | C48B6.2 | Embryo | NA |
| WBGene00016907 | C53H9.2 | Embryo | NA |
| WBGene00016977 | akap-1 | Embryo | NA |
| WBGene00016989 | mrpl-48 | Embryo | NA |
| WBGene00017159 | F01F1.2 | Embryo | NA |
| WBGene00017210 | F07E5.5 | Embryo | NA |
| WBGene00017319 | mrps-9 | Embryo | NA |
| WBGene00017356 | F10E9.4 | Embryo | NA |
| WBGene00017676 | F21F3.7 | Embryo | NA |
| WBGene00017763 | crls-1 | Embryo | NA |
| WBGene00017769 | hmgs-1 | Embryo | NA |
| WBGene00017776 | F25B5.5 | Embryo | NA |
| WBGene00017830 | rpb-8 | Embryo | NA |
| WBGene00017874 | acdh-9 | Embryo | NA |
| WBGene00017889 | F28B3.10 | Embryo | NA |
| WBGene00017924 | mrps-21 | Embryo | NA |
| WBGene00017959 | ugt-42 | Embryo | NA |

| | | | |
|---|---|---|---|
| WBGene00018046 | coa-3 | Embryo | NA |
| WBGene00018107 | F36H5.9 | Embryo | NA |
| WBGene00018240 | cox-17 | Embryo | NA |
| WBGene00018305 | exos-8 | Embryo | NA |
| WBGene00018339 | abcf-3 | Embryo | NA |
| WBGene00018422 | F44E2.9 | Embryo | NA |
| WBGene00018431 | F44E7.9 | Embryo | NA |
| WBGene00018475 | F45E12.5 | Embryo | NA |
| WBGene00018599 | F48C1.6 | Embryo | NA |
| WBGene00018674 | F52C9.3 | Embryo | NA |
| WBGene00018723 | F53A3.7 | Embryo | NA |
| WBGene00018762 | F53E10.6 | Embryo | NA |
| WBGene00018890 | F55F8.2 | Embryo | NA |
| WBGene00018891 | F55F8.3 | Embryo | NA |
| WBGene00018898 | F55F10.1 | Embryo | NA |
| WBGene00018934 | F56B3.11 | Embryo | NA |
| WBGene00018961 | mrps-16 | Embryo | NA |
| WBGene00019058 | F58F9.4 | Embryo | NA |
| WBGene00019076 | mrpl-24 | Embryo | NA |
| WBGene00019249 | hrpa-2 | Embryo | NA |
| WBGene00019275 | rpac-40 | Embryo | NA |
| WBGene00019294 | K02A6.3 | Embryo | NA |
| WBGene00019328 | clec-149 | Embryo | NA |
| WBGene00019380 | K04C2.2 | Embryo | NA |
| WBGene00019594 | K09H9.1 | Embryo | NA |
| WBGene00019658 | K11H12.1 | Embryo | NA |
| WBGene00019678 | K12H4.3 | Embryo | NA |
| WBGene00019800 | mtss-1 | Embryo | NA |
| WBGene00019812 | R01H2.4 | Embryo | NA |
| WBGene00019836 | R02F2.7 | Embryo | NA |
| WBGene00020036 | R12E2.11 | Embryo | NA |
| WBGene00020037 | mrps-6 | Embryo | NA |
| WBGene00020067 | clec-158 | Embryo | NA |
| WBGene00020089 | R119.3 | Embryo | NA |
| WBGene00020097 | larp-1 | Embryo | NA |
| WBGene00020171 | T02H6.1 | Embryo | NA |
| WBGene00020189 | tfbm-1 | Embryo | NA |
| WBGene00020269 | erfa-1 | Embryo | NA |
| WBGene00020273 | urb-1 | Embryo | NA |
| WBGene00020296 | rrp-8 | Embryo | NA |
| WBGene00020389 | T10B5.3 | Embryo | NA |
| WBGene00020717 | mrpl-4 | Embryo | NA |
| WBGene00020718 | mrps-2 | Embryo | NA |
| WBGene00020902 | jmjc-1 | Embryo | NA |
| WBGene00020993 | mtrf-1L | Embryo | NA |
| WBGene00021021 | mrpl-30 | Embryo | NA |
| WBGene00021026 | W04C9.4 | Embryo | NA |

| | | | |
|---|---|---|---|
| WBGene00021073 | nsun-1 | Embryo | NA |
| WBGene00021074 | gldi-1 | Embryo | NA |
| WBGene00021078 | unc-132 | Embryo | NA |
| WBGene00021276 | Y23H5B.5 | Embryo | NA |
| WBGene00021277 | ddx-10 | Embryo | NA |
| WBGene00021286 | tald-1 | Embryo | NA |
| WBGene00021377 | Y37E11B.5 | Embryo | NA |
| WBGene00021595 | Y46E12BL.2 | Embryo | NA |
| WBGene00021660 | nol-14 | Embryo | NA |
| WBGene00021697 | gcn-1 | Embryo | NA |
| WBGene00021715 | Y49F6B.2 | Embryo | NA |
| WBGene00021757 | taco-1 | Embryo | NA |
| WBGene00021829 | mrpl-17 | Embryo | NA |
| WBGene00021830 | Y54E10A.10 | Embryo | NA |
| WBGene00021843 | nobh-1 | Embryo | NA |
| WBGene00021899 | grwd-1 | Embryo | NA |
| WBGene00021920 | mrps-25 | Embryo | NA |
| WBGene00021930 | Y55F3AM.13 | Embryo | NA |
| WBGene00022016 | Y61A9LA.4 | Embryo | NA |
| WBGene00022046 | garr-1 | Embryo | NA |
| WBGene00022071 | spig-14 | Embryo | NA |
| WBGene00022072 | cpg-9 | Embryo | NA |
| WBGene00022107 | Y71F9AL.1 | Embryo | NA |
| WBGene00022148 | ddx-27 | Embryo | NA |
| WBGene00022159 | mppa-1 | Embryo | NA |
| WBGene00022275 | txt-7 | Embryo | NA |
| WBGene00022371 | set-29 | Embryo | NA |
| WBGene00022373 | mrpl-15 | Embryo | NA |
| WBGene00022378 | Y94H6A.5 | Embryo | NA |
| WBGene00022394 | natb-1 | Embryo | NA |
| WBGene00022470 | mrpl-19 | Embryo | NA |
| WBGene00022484 | fbxa-77 | Embryo | NA |
| WBGene00022500 | lfi-1 | Embryo | NA |
| WBGene00022538 | ZC190.4 | Embryo | NA |
| WBGene00022583 | mrps-18A | Embryo | NA |
| WBGene00022792 | ZK686.2 | Embryo | NA |
| WBGene00023068 | rpl-41.2 | Embryo | NA |
| WBGene00023419 | F53F4.16 | Embryo | NA |
| WBGene00023422 | mrpl-21 | Embryo | NA |
| WBGene00023487 | mrps-24 | Embryo | NA |
| WBGene00044026 | cisd-1 | Embryo | NA |
| WBGene00044094 | fitm-2 | Embryo | NA |
| WBGene00044318 | tag-267 | Embryo | NA |
| WBGene00044321 | mrps-30 | Embryo | NA |
| WBGene00044344 | mrpl-39 | Embryo | NA |
| WBGene00044440 | adpr-1 | Embryo | NA |
| WBGene00044511 | clec-181 | Embryo | NA |

| | | | |
|---|---|---|---|
| WBGene00044789 | T07A9.14 | Embryo | NA |
| WBGene00044894 | Y71F9AR.4 | Embryo | NA |
| WBGene00045433 | F49D11.10 | Embryo | NA |
| WBGene00050893 | Y94A7B.11 | Embryo | NA |
| WBGene00050913 | T12B5.14 | Embryo | NA |
| WBGene00050939 | C05G5.7 | Embryo | NA |
| WBGene00195010 | Y37E11AL.12 | Embryo | NA |
| WBGene00235258 | T05C7.5 | Embryo | NA |
| WBGene00005137 | srd-60 | Adult | NA |
| WBGene00006274 | str-247 | Adult | NA |
| WBGene00000177 | aqp-9 | Adult | NA |
| WBGene00000287 | cal-3 | Adult | NA |
| WBGene00000780 | cpn-4 | Adult | NA |
| WBGene00000988 | dhs-25 | Adult | NA |
| WBGene00001772 | gst-24 | Adult | NA |
| WBGene00002011 | hsp-12.2 | Adult | NA |
| WBGene00006125 | str-61 | Adult | NA |
| WBGene00006583 | tnc-2 | Adult | NA |
| WBGene00006586 | tni-4 | Adult | NA |
| WBGene00006588 | tnt-3 | Adult | NA |
| WBGene00006589 | tnt-4 | Adult | NA |
| WBGene00006825 | unc-96 | Adult | NA |
| WBGene00007122 | B0250.5 | Adult | NA |
| WBGene00007240 | C01G10.15 | Adult | NA |
| WBGene00008294 | clec-11 | Adult | NA |
| WBGene00008869 | F15G9.6 | Adult | NA |
| WBGene00009436 | marb-1 | Adult | NA |
| WBGene00009475 | F36F2.1 | Adult | NA |
| WBGene00009730 | myo-6 | Adult | NA |
| WBGene00011270 | R31.2 | Adult | NA |
| WBGene00011573 | anmt-3 | Adult | NA |
| WBGene00012158 | ucr-2.1 | Adult | NA |
| WBGene00012726 | Y39G8B.5 | Adult | NA |
| WBGene00013237 | ndab-1 | Adult | NA |
| WBGene00013620 | fbxa-111 | Adult | NA |
| WBGene00014052 | ZK669.2 | Adult | NA |
| WBGene00014063 | clec-143 | Adult | NA |
| WBGene00014088 | ZK809.8 | Adult | NA |
| WBGene00015142 | B0310.6 | Adult | NA |
| WBGene00015723 | gtnt-6 | Adult | NA |
| WBGene00015837 | math-8 | Adult | NA |
| WBGene00016668 | ilys-1 | Adult | NA |
| WBGene00016898 | clik-2 | Adult | NA |
| WBGene00017426 | F13C5.5 | Adult | NA |
| WBGene00018388 | F43C11.11 | Adult | NA |
| WBGene00019361 | clik-3 | Adult | NA |
| WBGene00019661 | K11H12.5 | Adult | NA |

| | | | |
|---|---|---|---|
| WBGene00019818 | R02C2.7 | Adult | NA |
| WBGene00019937 | R07E4.3 | Adult | NA |
| WBGene00020192 | T03F1.11 | Adult | NA |
| WBGene00021031 | W04H10.2 | Adult | NA |
| WBGene00022075 | nduv-3 | Adult | NA |
| WBGene00022615 | ZC449.5 | Adult | NA |
| WBGene00023420 | C26D10.7 | Adult | NA |
| WBGene00044260 | Y87G2A.19 | Adult | NA |
| WBGene00044413 | B0205.12 | Adult | NA |
| WBGene00044427 | F56A6.5 | Adult | NA |
| WBGene00044754 | Y119C1B.12 | Adult | NA |
| WBGene00045483 | oxy-5 | Adult | NA |
| WBGene00195212 | F09B12.7 | Adult | NA |
