## Supplementary material for "Confidence-aware learning for transcriptome-based prediction of OXPHOS genes in *Caenorhabditis elegans* under incomplete functional annotation": Table S7

| Algorithm | Hyperparameter | Range |
| --- | --- | --- |
| **RF** | n_estimators | np.arange(1, 20, 1) |
| | max_depth | np.arange(1, 20, 1) |
| | min_samples_split | np.arange(1, 20, 1) |
| | min_samples_leaf | np.arange(1, 20, 1) |
| **SVM** | kernel | linear', 'poly', 'rbf', 'sigmoid' |
| | C | np.arange(0.1, 20, 1) |
| | gamma | np.arange(0.01, 10, 0.5) |
| **KNN** | n_neighbors | np.arange(2, 30, 2) |
| | weights | uniform', 'distance' |
| | algorithm | auto', 'ball_tree', 'kd_tree', 'brute' |
| | leaf_size | np.arange(1, 100, 10) |
